# IDSL.CSA: Composite Spectra Analysis for Chemical Annotation of Untargeted Metabolomics Datasets

**DOI:** 10.1101/2023.02.09.527886

**Authors:** Sadjad Fakouri Baygi, Yashwant Kumar, Dinesh Kumar Barupal

**Author notes:** Corresponding author: Address: CAM Building, 3rd floor, 17 E 102nd St, New York, NY 10029.

## Abstract

Poor chemical annotation of high-resolution mass spectrometry data limit applications of untargeted metabolomics datasets. Our new software, the Integrated Data Science Laboratory for Metabolomics and Exposomics – Composite Spectra Analysis (IDSL.CSA) R package, generates composite mass spectra libraries from MS1-only data, enabling the chemical annotation of LC/HRMS peaks regardless of the availability of MS2 fragmentation spectra. We demonstrate comparable annotation rates for commonly detected endogenous metabolites in human blood samples using IDSL.CSA libraries versus MS/MS libraries in validation tests. IDSL.CSA can create and search composite spectra libraries from any untargeted metabolomics dataset generated using high-resolution mass spectrometry coupled to liquid or gas chromatography instruments. The cross-applicability of these libraries across independent studies may provide access to new biological insights that may be missed due to the lack of MS2 fragmentation data. The IDSL.CSA package is available in the R CRAN repository at https://cran.r-project.org/package=IDSL.CSA. Detailed documentation and tutorials are provided at https://github.com/idslme/IDSL.CSA.

**For Table of Contents Only:** 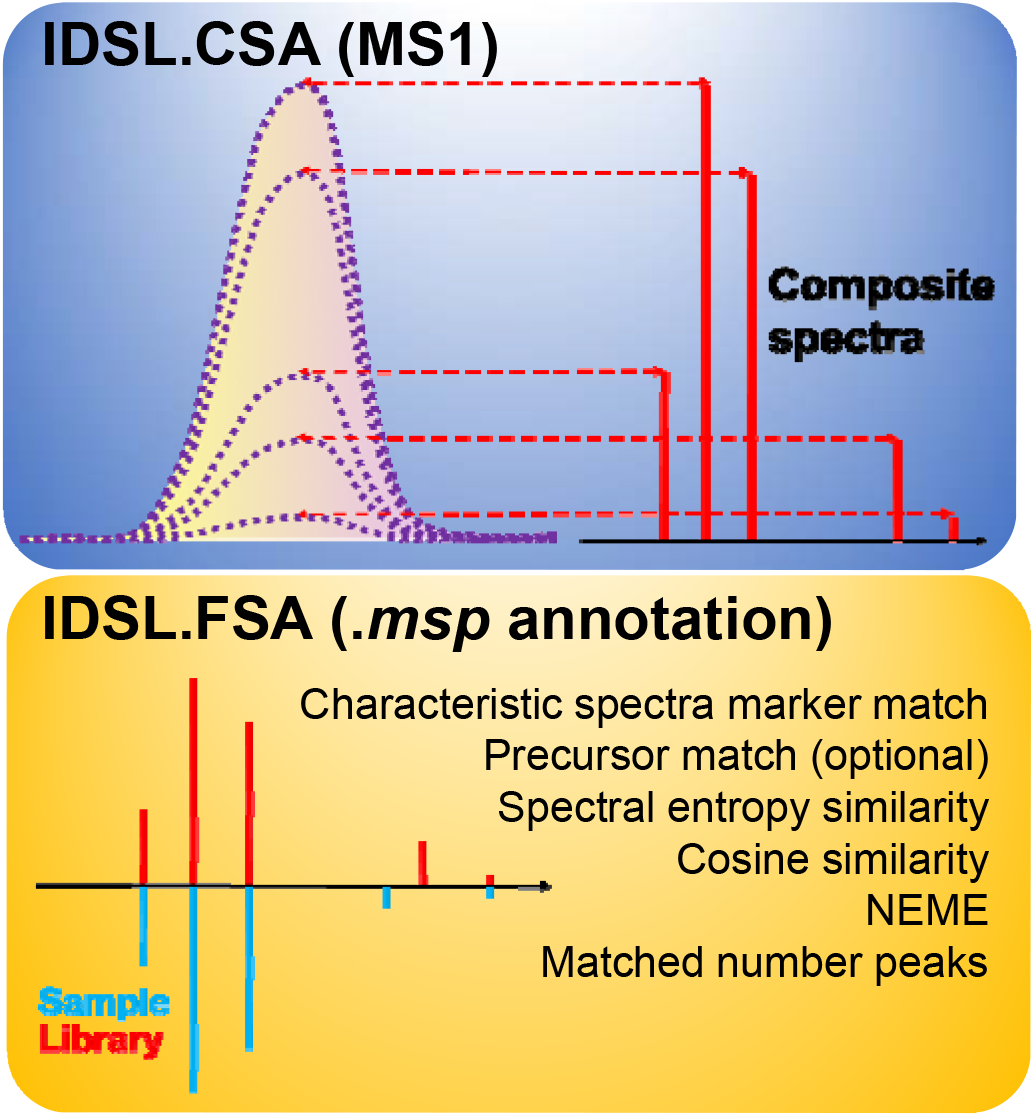

## Introduction

Metabolomics has a great potential to uncover biological mechanisms and biomarkers that can be translated to innovative products to improve human health, food security and the environment. One major approach of metabolomics is the untargeted profiling of the small molecule chemical space in a biospecimen to create large databases of chemical measurements under different phenotype^1^ and genotype backgrounds^1, 2^ . High-resolution mass spectrometry coupled with liquid chromatography (LC/HRMS) is the most commonly used technique for untargeted metabolomics assays. However, not all peaks in an LC/HRMS dataset are annotated with chemical information, mainly for two reasons 1) MS2 fragmentation data are not available for every detected LC/HRMS peak and 2) Available MS2 libraries poorly cover the expected chemical space. As a result, a large volume of untargeted metabolomics data remains underutilized, leading to missed discovery opportunities.

A single chemical compound can generate several ions species due to chemical reactions including fragmentation and adduct formations in the electrospray ionization (ESI) source, collectively known as the MS1 data. Sporadically, selected ions (precursors) can be fragmented further to generate mass spectra (MS2) that can be searched against MS/MS mass spectral libraries such as NIST 2020 for chemical annotation^3, 4^ along with several advances in annotating and interpreting MS/MS data^5-7^ . Existing approaches^8-11^ for MS1 data annotation are limited to group the correlating ions in order to flag LC/HRMS peaks as potential isotopologues, adducts or in-source fragments or assigning molecular formula using isotope patterns^12^. Promisingly, these approaches including RAMClust^8^, eISA^11^, CorrDec^13^, IIMN^10^, MS-FLO^14^, and MetaboAnnotatoR^15^ also suggests that co-eluting ions at MS1 level can be exported as a spectra which may match an entry in a MS/MS library, enabling the chemical annotation of untargeted metabolomics data using only MS1 data. Consistent ion annotation for common metabolites across multiple studies suggests that related ESI signals for a compound are trackable and can be utilized to annotate LC/HRMS peaks^12, 16-19^ . There is a need to advance computational resources to annotate compounds using these grouped peaks in untargeted LC/HRMS datasets. Converting these correlating ions on MS1 level to re-usable mass spectra libraries can be useful for annotating peaks in independent studies that may or may not have MS2 data.

Here, we propose to create composite spectra libraries of correlating LC/HRMS peaks in the MS1-only data from untargeted metabolomics assays and to use these libraries for annotating peaks by mass spectral library searches. For that, we have developed a new R package IDSL.CSA (https://cran.r-project.org/package=IDSL.CSA) for creating CSA libraries and a companion R package IDSL.FSA (https://cran.r-project.org/package=IDSL.FSA) for mass spectral similarity searches.

## Material and methods

### Publicly available LC/HRMS test datasets

Tables S.1 presented the mass spectrometry data and known annotations used in this work. MSV000088661, ST000923 and ST001000 from Massive UCSD (https://massive.ucsd.edu), Metabolomics WorkBench (https://www.metabolomicsworkbench.org) data repositories. Raw mass spectrometry data were converted to the centroid mzML format using the MSConvert utility of ProteoWizard.

### R packages

IDSL.CSA R package (version 1.0) has been provided via the R-CRAN repository (https://cran.r-project.org/package=IDSL.CSA). IDSL.MXP R package^19^ (https://CRAN.R-project.org/package=IDSL.MXP) was used to read LC/HRMS files in mzML/mzXML/netCDF data formats. IDSL.IPA R package (version 2.6)^19^ (https://CRAN.R-project.org/package=IDSL.IPA) was used for generating chromatographic peak lists and aligned peak tables for each study. IDSL.FSA R package (version 1.0) provided at (https://cran.r-project.org/package=IDSL.FSA) was used for mass spectral similarity searches.

### IDSL.CSA workflow

Initially, mzML/mzXML/netCDF mass spectrometry data were processed using the IDSL.IPA package^19^ to generate individual peak lists, aligned peak tables and the intra-sample correlations among all detected peaks for a study. For IDSL.CSA data processing, IDSL.IPA provides essential information including peak boundaries and ^12^ C/^13^ C ion pair values for the detected peaks. A chromatograph peak is a region of an extracted ion chromatogram for ^12^ C ion which is also accompanied with frequent ^13^ C ion pair values. To present a comprehensive annotation and deconvolution data processing workflow IDSL.CSA workflow encapsulate various modules to generate three types of mass spectral libraries – 1) composite spectra (CSA) 2) data-dependent acquisition (DDA) and 3) data independent acquisition (DIA).

**CSA** peak deconvolution method (Figure 1) detects MS1 chromatographic peaks potentially originated from the same compound (Figure 2). The method initially sorts the all MS1 ^12^ C peaks from an IDSL.IPA peak list by their intensity, and the most intense peak at a user-provided signal-to-noise (S/N) ratio threshold is selected as the seed peak for creating CSA spectra. Each peak was assigned to only a single CSA spectrum to avoid redundancies. Only peaks correlating on the aligned peak height table were selected to remove false positives (Section S.1). A retention time window is then used to group potentially co-eluting ions available in the IDSL.IPA peak list. Then we extracted EICs for each ion. The Pearson correlation is computed among EICs of grouped peaks after a LOESS smoothing followed by a cubic spline interpolation. The correlations were calculated only for data points above a height percentage of chromatographic peaks to minimize effects of peak tailings and frontings. To avoid CSA spectra duplication, each ^12^ C peak is assigned to only one CSA spectra. Thresholds for the minimum number of the CSA ions and the minimum distance between lowest and highest ions were used to prevent only collecting isotopic envelopes and low-quality CSA spectra such as clusters of only two ions. ^13^ C ions are streamlined for CSA analysis only when their ^12^ C counterpart was matched in a CSA cluster. All CSA spectra are deconvoluted first for each sample. Then the user-provided m/z value with retention time was checked to link the CSA spectra to a true positive annotation (if available). **DDA:** DDA spectra with precursor details were extracted from mass spectrometry data (mzML). When more than one DDA spectra with the same precursor m/z were observed within a boundary of an IDSL.IPA peak, three options were provided 1) to integrate all DDA spectra with the same precursor 2) to select the most abundant 3) to de-noise spectra by a applying a Pearson correlation coefficient threshold after removing flat peaks using a relative standard deviations threshold for the DDA spectra. **DIA:** DIA spectra were created by the same approach used for CSA, except that the fragment ions were obtained from unprocessed m/z values at MS level = 2 (MS^e^ or SWATH-MS) data channels from the nearest MS2 scan to the MS1 scan at the apex of the MS1 chromatographic peak.^16^ DIA method uses the MS1 peak as the precursor peak. Comprehensive tutorials on parameter selection and workflow for these methods is provided in the Wiki tab of the IDSL.CSA package at https://github.com/idslme/IDSL.CSA/wiki. Spectra from these three methods were exported to the MSP format. Parameter files are provided at https://zenodo.org/record/7530387.

**Figure 1.**
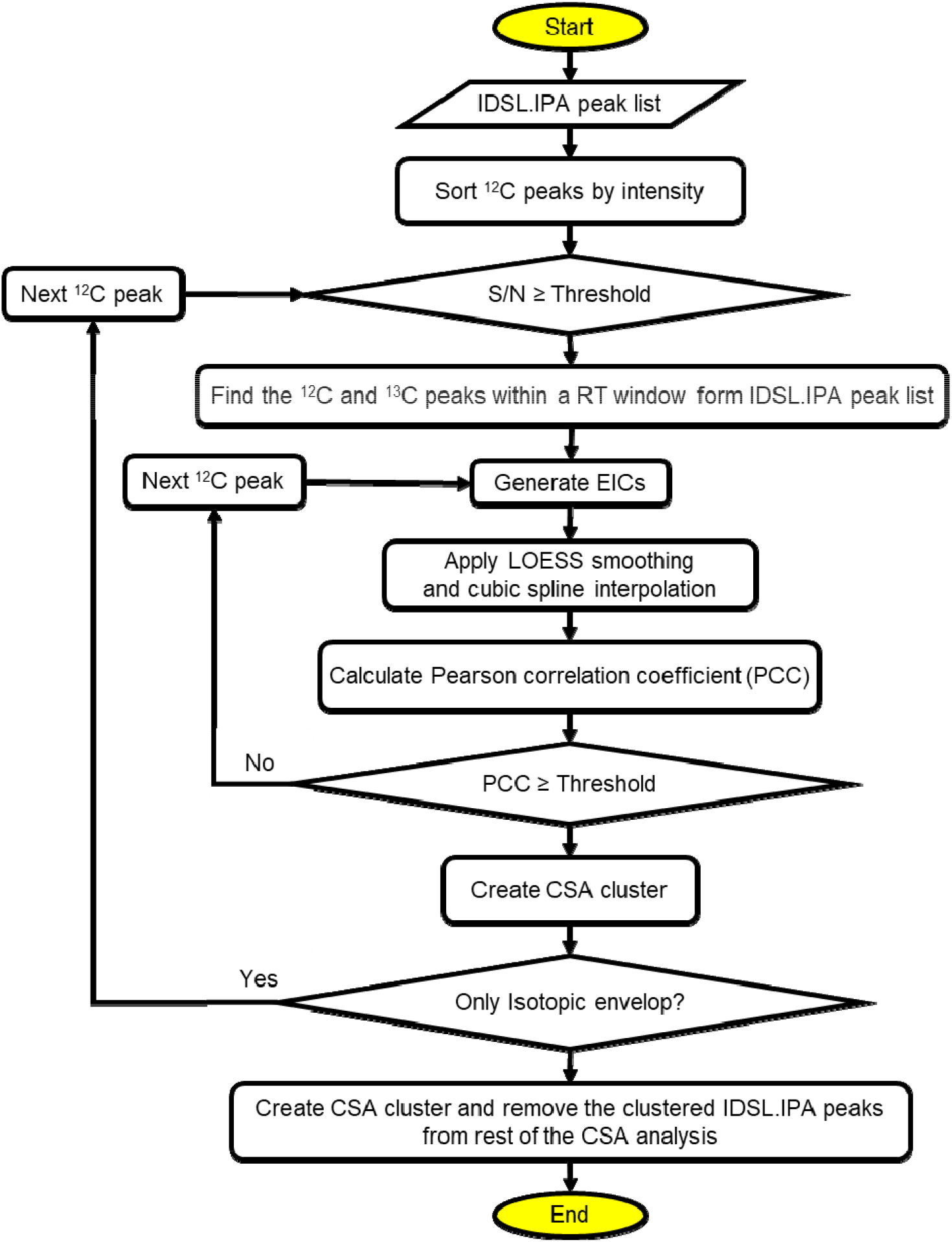
Flowchart of the Composite Spectra Analysis workflow.

**Figure 2.**
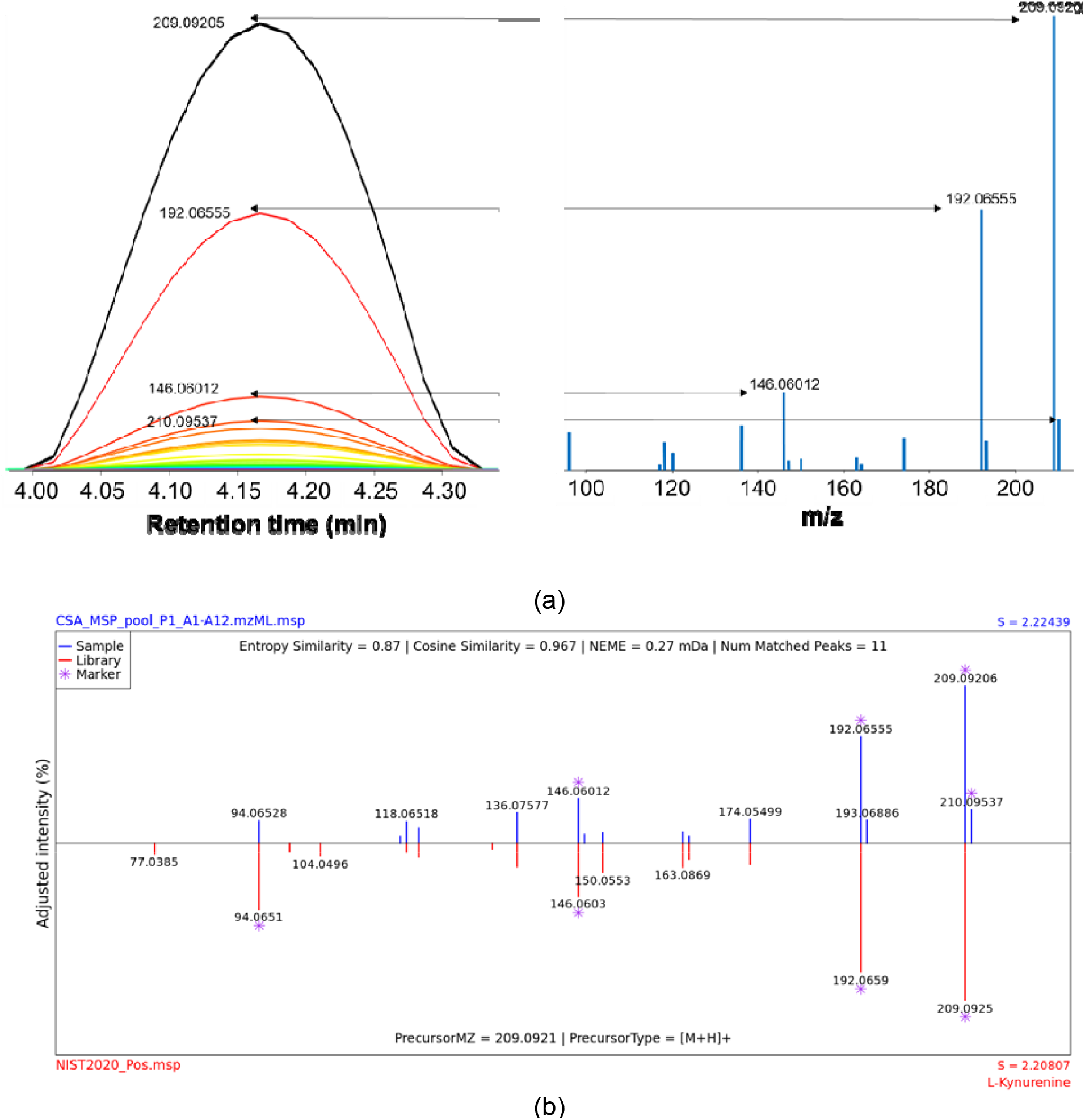
a) Schematic of composite spectra creation from aligned extracted chromatograms in MSV000088661 (IROA standards) study for Kynurenine by the IDSL.CSA package. b) Annotating the CSA spectra from panel (a) using the Kynurenine NIST reference MS/MS spectra by IDSL.FSA package (noise removal = 1%).

### Unique spectra detection and aggregation

IDSL.CSA can aggregate similar CSA/DDA/DIA spectra from targeted msp library generation and untargeted screening workflows in the batch studies with many numbers of samples. By aggregating similar spectra, IDSL.CSA can isolate unique spectra with high frequency of detection and reduce the complexity of the metabolomics data to lower the load of identification workflows. Only for the CSA workflow, unique spectra aggregation also can be performed on the aligned peak tables from IDSL.IPA to generate CSA spectra with the most common ions across the entire batch using Tanimato coefficient thresholds.

To perform mass spectral similarity search, IDSL.FSA package (https://CRAN.R-project.org/package=IDSL.FSA) was developed to annotate CSA spectra that do not include precursor values in addition to DDA and DIA exported msp files. IDSL.FSA is able to annotate all kinds of msp and mgf file using intelligent pre-filtering steps (see Section S.2). Parameter selection is doable through using a Microsoft excel format (https://zenodo.org/record/7530387) without prior library curation. A complete tutorial on IDSL.FSA is provided at https://github.com/idslme/IDSL.FSA. If mass spectral libraries for standards were from different sources and in different formats such as mgf, they were converted to a standard msp file and separated by ionization mode (+/-). To enable faster searches in R, msp files for reference compounds and authentic standards were converted into a fragmentation spectra database (FSDB) that stored pre-calculated spectral entropy for each spectra in the MSP file (https://zenodo.org/record/7530387). FSDBs for public databases including the Global Natural Product Social Molecular Networking (GNPS) and Mass bank of North America (MoNA) for positive and negative modes are provided at (https://zenodo.org/record/7530397). Similar to Li *et. al*.,^20^ adjacent fragments within the instrument resolution were resolved followed by a noise removal (Default = 1%). A threshold above a baseline (%) was used to determine characteristic spectra markers, and then a minimum threshold (%) of matched characteristic spectra markers was used to allow partial symmetry matches in case of presence of instrumental noises. Cosine and entropy similarity scores were computed for spectra matching. The cosine similarity was computed using equation (1) only using the library fragments.

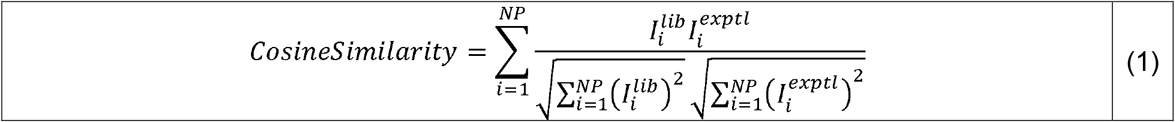

where *I*_*i*_ and *NP* represent the intensity of the fragment, and number of matched fragment peaks in the fragmentation spectra, respectively. Superscripts of *lib* and *exptl* represent library and experimental fragmentation spectra, respectively. Entropy similarity was calculated from spectral entropy (*S*) values described by Li *et. al*.^20^

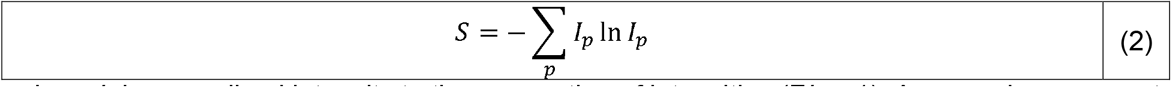

where *I*_*p*_ is normalized intensity to the summation of intensities (Σ*I*_*p*_ = 1). A merged mass spectra *lib:exptl* is generated by 1:1 mixing the normalized *lib* and *exptl* spectra. Then, the entropy similarity was calculated using equation (3).

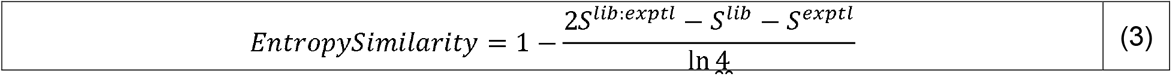

An additional weight transformation proposed by Li *et. al*.,^20^ was applied to adjust spectral entropy using equation (4).

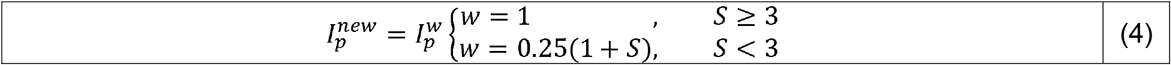

The transformed 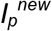 should also be re-normalized 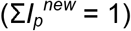. An entropy similarity cutoff (Default ≥ 0.75) may be used to filter out candidate hits. Normalized Euclidean mass error (*NEME*) was calculated using the equation (5) to further asses the fragmentation spectra.

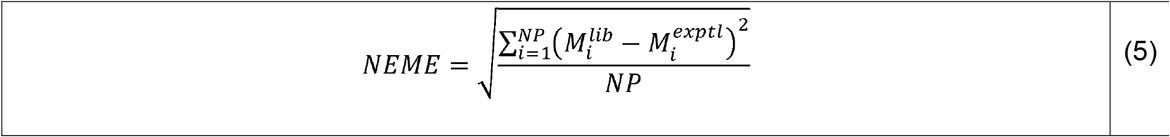

where *M*_*i*_ represents the mass of the fragmentation peaks. A maximum threshold for *NEME* may be used to cut off hits with higher mass errors.

When the entire candidate hits were matched for an experimental fragmentation spectrum, the candidate hits are sorted using the following equation.

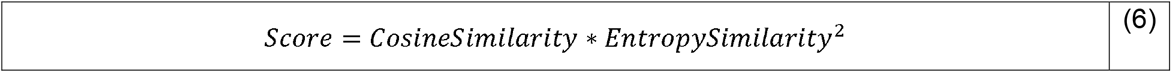

For each sample in a study, spectra search result table was generated. An example table is provided at (https://zenodo.org/record/7530263). If the aligned table generated by the IDSL.IPA R package was available, spectra search results were summarized for annotation frequency and overall median ranks.

### Parameter files

IDSL.IPA and IDSL.CSA parameter files for MSV000088661, ST000923 and ST001000 studies are provided at (https://zenodo.org/record/7530409). These files include – 1) creating reference CSA spectra for authentic standard 2) creating CSA spectra for true positive annotation ST001000 study and 3) creating CSA spectra in a full deconvolution mode for ST001000 and 4) IDSL.IPA data processing settings. After filling up parameters in these files, the excel sheets were used directly as an input for running the workflow function in the IDSL.CSA R package.

### Statistical analysis

Student *t*-test and ChemRICH analysis was performed in R to find significantly different chemical and chemical sets in the Crohn’s disease patients in comparison to the healthy control group.

## Results

First, we analyzed the elution profile of Kynurenine reference compounds in LC/HRMS data by the IDSL.IPA software^19^ for a test file (pool_P1_A1-A12.mzML) from the MSV000088661 study. IDSL.IPA detected 255 chromatographic peaks (intensity threshold ≥ 10^5^ and S/N ≥ 5) for this test file of a single mixture of 12 authentic standards (https://zenodo.org/record/7530009). Out of those, 153 (60%) had at least one neighboring peak in a retention time window of 0.01 minutes. Many of these co-occurring ions may have originated from a single compound eluting from the LC column. For example, a cluster of ions (209.292, 192.065, 146.060, …) related to Kynurenine compound standard followed an almost identical elution profile (Figure 2) at retention time 4.18 minutes which also matched Kynurenine reference spectra from NIST MS/MS library. To automatically test if those neighboring peaks are related to a single compound, we have developed a new spectra deconvolution algorithm to group them by measuring the elution profile similarity. Extracted ion chromatograms (EICs) for individual peaks were extracted from the raw data file. Then, EICs were smoothed using the Local Polynomial Regression Fitting (LOESS) regression to provide a robust estimate of the elution profile similarity. At a Pearson correlation coefficient threshold of 0.98 between the EICs, 76.9% peaks were grouped into 42 EIC groups or CSA clusters (https://zenodo.org/record/7530028) for the test file. Clusters were selected only if they had at least two peaks from the IDSL.IPA peak list (https://zenodo.org/record/7530009) and a difference between minimum and maximum m/z values was greater than eight. These two criteria ensured that the CSA clusters had m/z values outside the isotopologue ranges which obviously have highly similar elution profiles. Observations of these clusters motivated us to explore their utility in annotating compounds in untargeted LC/HRMS datasets.

We checked if a CSA cluster can represent a deconvoluted mass spectra for a compound. For each CSA cluster, we exported all m/z values within a cluster, the retention time at the apex, the peak height values at the apex of for each ion’s EIC as the intensity values, the correlation statistics and the additional metadata to a mass spectra file in the NIST MSP format (https://zenodo.org/record/7530023). The file had 42 spectra. They were named ‘CSA Spectra’. Searching these spectra against the NIST 2020 library suggested high confidence annotations for 5 clusters (https://zenodo.org/record/7530063). See the method section for spectral search parameters. Top hits for these clusters were Purine, L-Kynurenine, N-Acetyl histidine, Homoserine, N-Acetylneuraminic acid (Figure S.1). The annotations were also confirmed by the MS/MS spectra (Figure S.2, https://zenodo.org/record/7530151). These results suggested that CSA Spectra may be used for annotating compounds in LC-HRMS data with limited or no availability of MS/MS data which is consistent with previous reports (cite RAMClust^8^ and MetaboAnnotatorR^15^).

We developed a workflow for automatically creating a CSA spectra library for authentic standards. A total of 359 unique chemical standards had confirmed reverse phase retention time and precursor m/z values across 54 LC/HRMS data in the ESI positive mode (https://zenodo.org/record/7530170). For these standards, 214(59.6%) were associated with a CSA spectrum by the workflow. Only up to 42(19.6%) CSA spectra contain similar in-source fragments to those of MS/MS spectra in reference libraries suggested that a majority of CSA spectra contained additional in-source ESI adducts that we do not expect for a MS/MS spectra. The spectra library is provided at (https://zenodo.org/record/7530184). The MTBLS1040^21^ dataset (https://zenodo.org/record/7879353), containing 124 standards analyzed at seven concentrations was processed to test the effect of concentration on CSA spectra creation. Only at the very low concentration levels, the number of ions is below >3, suggesting that low abundant compounds in from a sample may not have a CSA spectrum (Figure 3). We studied matrix effects on CSA spectra by comparing authentic standard compounds in study MSV000088661 and blood samples in study ST002044 that were processed using the same instrument and analytical conditions. Figure 4 indicated only 6 metabolites (out of 29) had CSA spectra similar to standard compounds in more than 50% MS1 peaks across 286 samples indicating the CSA spectra may differ by sample matrix types. Figure 4 also indicated that in general in 57% cases MS1 peaks cannot be clustered into CSA spectra in samples with matrix effect. We compared CSA spectra created for a BioRec human plasma sample analyzed in the same lab by Agilent QToF (ST001843) and Thermo Orbitrap (ST001264) instruments using the same chromatography gradient. A total of 337 CSA spectra were detected by both instruments and 69 were common among them (Figure 5).

**Figure 3.**
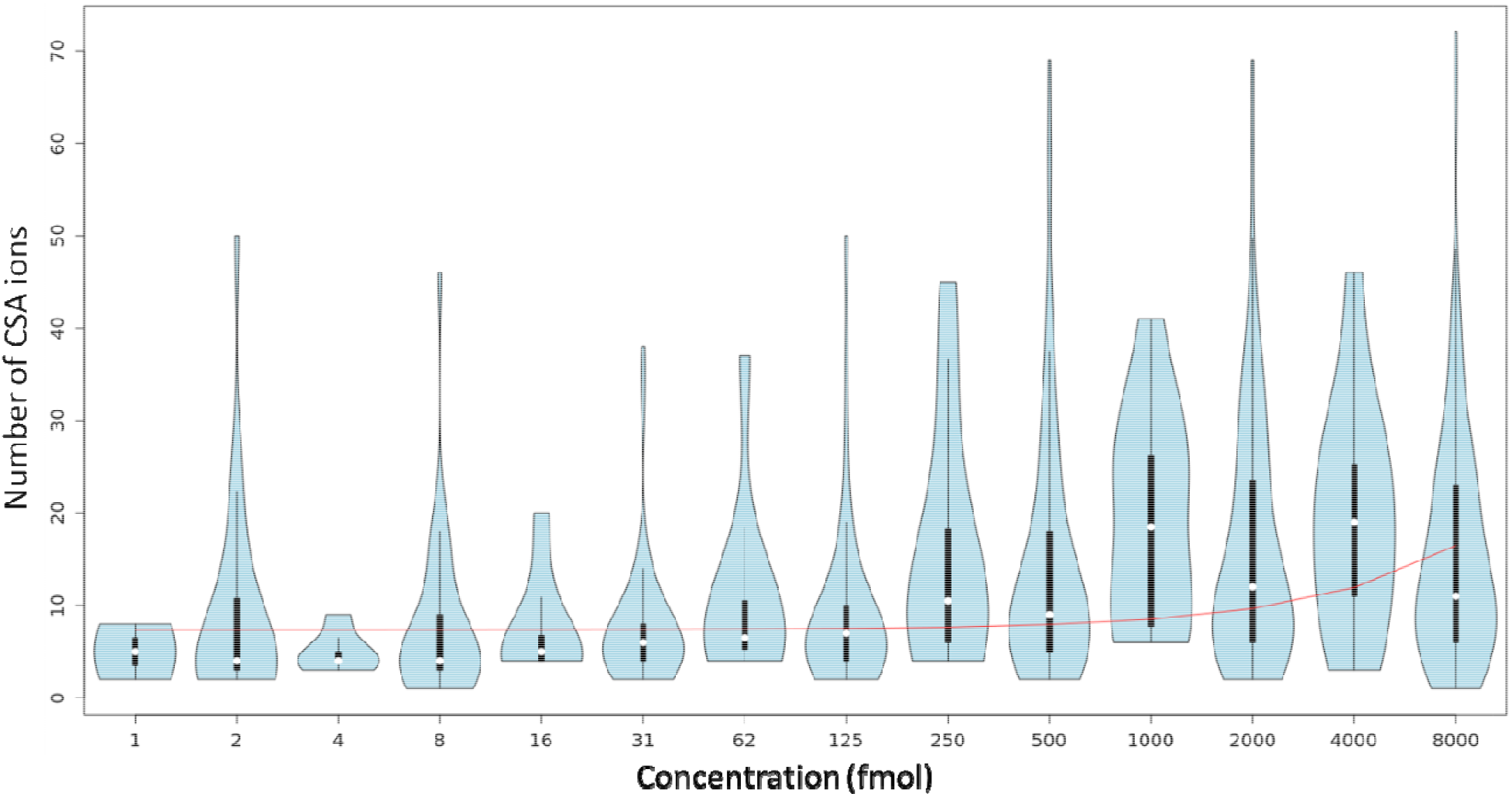
Effects of concentration on the number of ions in a CSA cluster for 124 neat standard compounds in MTBLS1040 study (https://zenodo.org/record/7879353).

**Figure 4.**
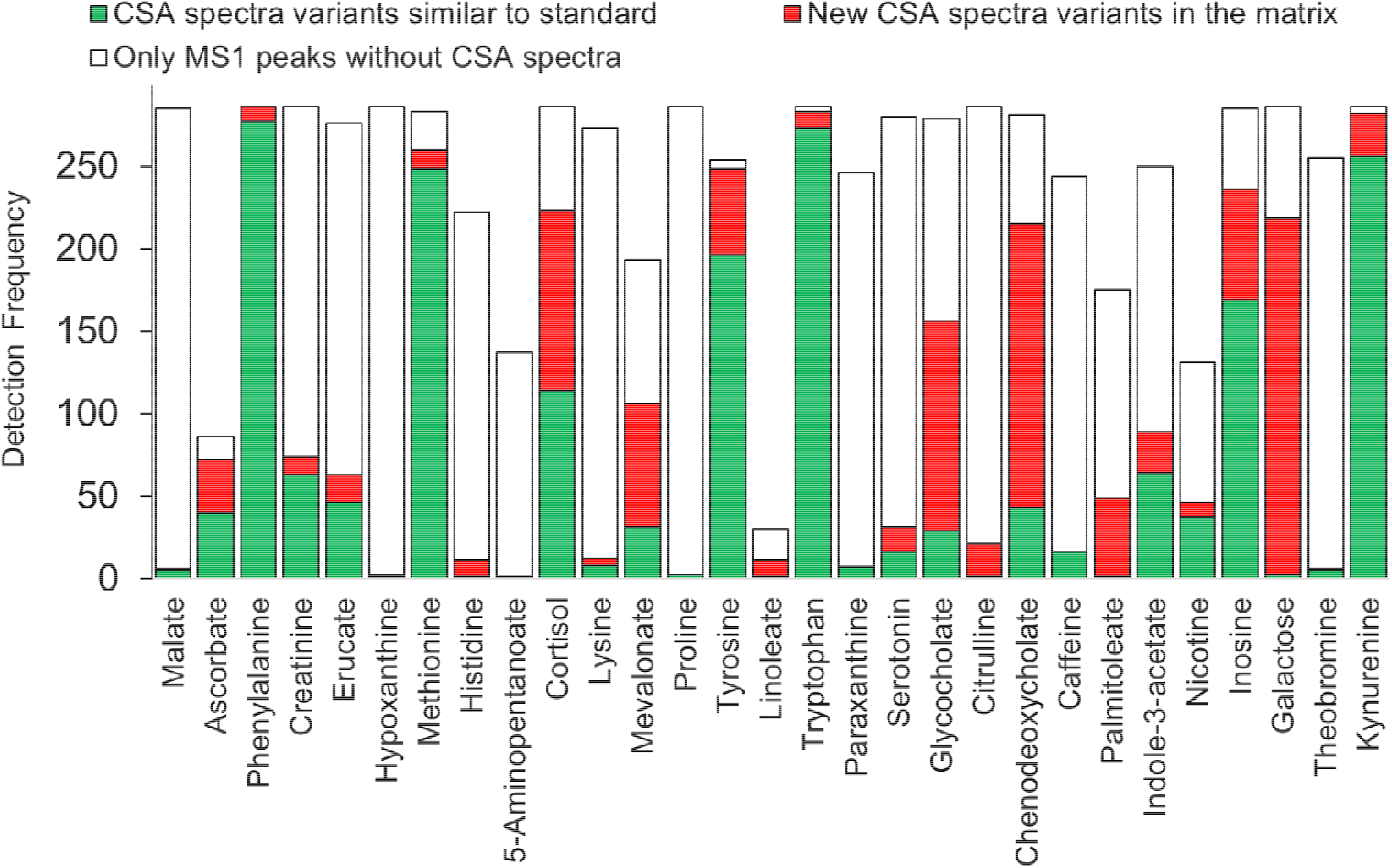
Matrix effects on CSA spectra variation for 29 metabolites across 286 samples in the ST002044 study.

**Figure 5.**
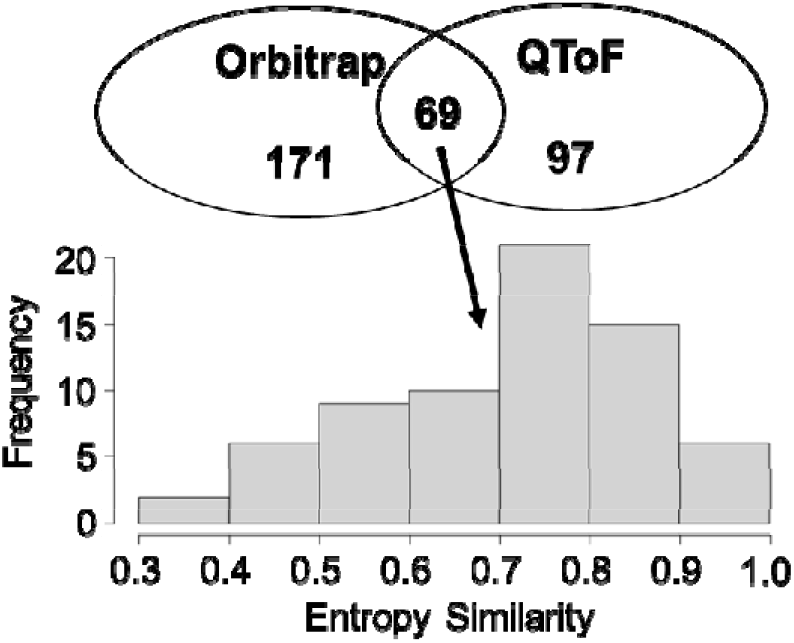
Comparison between Orbitrap (ST001264) and QToF (ST001843) instruments on a plasma sample analyzed for a lipidomic assay using samiliar chromatography methods.

The CSA workflow was extended to cover known annotations that have been reported for LC/HRMS data of biological specimens. In a publicly available untargeted metabolomics study (ST000923) of human stool samples, 177 annotations were targeted in 600 LC/HRMS data files (https://zenodo.org/record/7530227) for creating a CSA spectra library. Only MS1 data were available for this study. Up to 105(59.3%) compounds had an associated CSA spectra with a median frequency of 204, min of 1, max of 589 and a total of 25,351 CSA spectra were deconvoluted for the target compounds from 600 samples. Then, we identified variants of CSA spectra that are detected across multiple LC/HRMS data files. By using the entropy similarity approach^20^ (See method), we found that the average number of CSA spectra was 2.5 per compound. Searching these CSA spectra against mass spectral databases suggested that only 46(26%) had a hit in these libraries. This comparison highlights that a CSA spectrum can capture the in-source fragmentation that may resemble a known MS/MS spectrum, suggesting that the existing MS/MS libraries can also be used for annotating a subset of CSA spectra. The CSA spectra library for the annotated compounds for the study has been provided at (https://zenodo.org/record/7530237).

Next, we scaled and applied the CSA deconvolution algorithm to the ST001000 (n=226) study to detect all possible CSA spectra in LC/HRMS data collected for this study. Study data were collected using the HILIC chromatography and the ESI (+) mode ionization using a Thermo Q-Exactive Orbitrap mass spectrometry instrument. Only MS1 data was available for this study. For this study, a total of 139,018 CSA spectra were deconvoluted with a median of 631 per file. On average, 31% peaks from the MS1 peaklist were part of CSA spectra across all the samples. A total of 16,659 unique CSA spectra with at least 1% detection frequency were detected using a retention time window of 0.1 minute and an entropy similar threshold of 0.75 for this study (https://zenodo.org/record/7530245). Only 51 spectra were found in more than 90% samples in the study and 1/3 were detected in less than 5% of samples.

To annotate the deconvoluted CSA spectra, we have first searched them against MS/MS databases. For the ST001000 study, 802(4.8%) CSA spectra had a high confidence spectral similarity match (https://zenodo.org/record/7530263). Because a compound can have multiple CSA variants across the entire study, unique base peaks by chemical name and InChIKeys were selected to subset the aligned peak table for the study. A total of 377 unique peaks were selected and annotated with 321 unique InChIKeys and 343 unique chemical names (https://zenodo.org/record/7530275). This subset aligned table represents a data matrix for annotated compounds by the CSA workflow.

Next, we tested if a CSA spectra library created for one study can enable annotations of compounds in a different study. We searched the unique CSA spectra variants for ST001000 study against the CSA library that we have generated earlier for the study ST000923. The spectral search results (https://zenodo.org/record/7530302) suggested that 34 additional compounds (InChIKeys) can be annotated for study ST001000 that were not covered by the MS/MS library search (https://zenodo.org/record/7530310).

To verify the accuracy of the annotations obtained using only the spectral searches for CSA spectra, we have matched their retention time against the published data dictionaries for ST001000 and ST000923 studies (See methods). Data for both studies were collected by the same laboratory using identical analytical conditions, but ST000923 had 174 annotated compounds (InChIKeys), of which 89 were not previously annotated for the ST001000 study. Of the 488 annotations (InChIKeys) by CSA spectra, 60 were confirmed by matching their retention time in the published data dictionaries for ST000923 and ST001000 studies in the Metabolomics WorkBench repository. Only 26(26.4%) of the hits were probable false positives or in-source fragments. The IDSL.CSA workflow suggested 301 new annotations (unique InChIKeys) for the ST001000 study (https://zenodo.org/record/7530359). In this study, 7/42 (16%) compounds had a CSA spectrum at a different retention than the expected one for the true positive annotation. It can be suggested that the CSA creation had a 16% false positive rate.

To demonstrate the biological significance of the annotated compounds for the ST001000 study, we conducted a chemical set enrichment analysis using ChemRICH software^22^ . ChemRICH identified the chemical classes that were found to be significantly different between individuals with Crohn’s disease and healthy controls (https://zenodo.org/record/7530364). Our analysis revealed that several chemical classes, including cholic acids, amino acids, aminosalicylic acids, biogenic amines, hexosamines, and vitamins, were significantly different between these two groups (https://zenodo.org/record/7530366). The ChemRICH result suggests that the altered gut microbiota ecology^23^ in Crohn’s disease patients may also be connected to metabolic pathways involving these chemical classes.

Finally, we have incorporated the workflows for creating CSA libraries and annotating those using mass spectral similarity searches into two standard R package called ‘IDSL.CSA’ and ‘IDSL.FSA’ available on CRAN repository at (https://cran.r-project.org/package=IDSL.CSA) and. (https://cran.r-project.org/package=IDSL.FSA). We have also added workflows for processing data-dependent (DDA) and data-independent (DIA) acquisitions. The package includes a user-friendly parameter file in Microsoft Excel format that allows users to run different workflows and ensure reproducible data processing. Additionally, we also extended the CSA deconvolution to cover nominal mass data using a secondary ‘IDSL.NPA’ R package (https://cran.r-project.org/package=IDSL.NPA). The parameter tables are extensive and cover commonly used settings as well as several new parameters to optimize spectra deconvolution and spectral searches (Table S.2-5). Documentation, tutorials, and code for the software are available for IDSL.CSA and IDSL.FSA R packages in the GitHub repository at https://github.com/idslme/IDSL.CSA and https://github.com/idslme/IDSL.FSA, respectively.

## Discussion

We have developed a simple and easy to use integrated workflow of IDSL.CSA and IDSL.FSA R packages for annotating LC/HRMS peaks in untargeted metabolomics datasets using only MS1 data. The integrated approach using IDSL.IPA^19^, IDSL.CSA and IDSL.FSA R packages (Figure 6) contains easy to use steps for 1) creating CSA, DDA and DIA spectral libraries 2) performing mass spectral similarity searches using spectral entropy^20^, cosine similarity, normalized Euclidean mass error^12^ 3) prefiltering library spectra for faster searches (Section S.2) 4) refining deconvolution results aligned table (Section S.1) and 5) ranking annotations using spectra search results from all the samples within a study.

**Figure 6.**
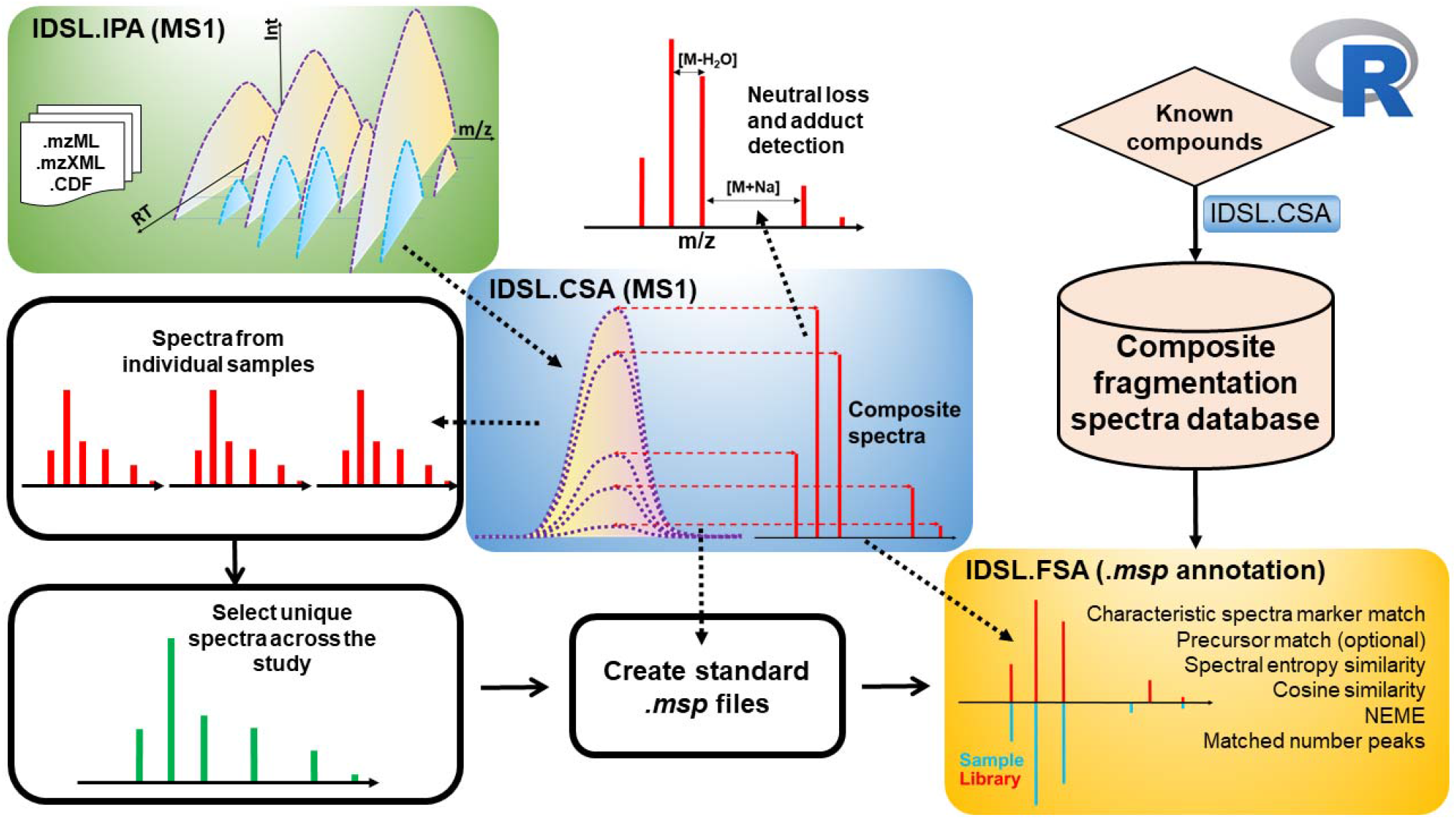
Integrated workflow of the IDSL.IPA, IDS.CSA and IDSL.FSA packages to deconvolute and annotate composite spectra.

We have developed a new workflow to create composite spectra libraries from LC-HRMS datasets. A CSA spectrum may be similar to a counter MS/MS spectrum of the compound but may often include additional ionization reactions (Figure S.3). As recommended by earlier reports^12, 16-19^, IDSL.CSA workflows also utilize both individual file level geometry alignment of LC/HRMS peaks and the co-detection frequency across all samples within a study to capture different variants of CSA spectra for a compound. For example, two variants of CSA spectra for kynurenine can contain different ESI adducts such as [M+Na]^+^ and [M-H_2_O+H]^+^ (Figure S.4). The composition of CSA variant spectra for a chemical depends on the instrument type, analytical method, sample matrix, gradient additives and other factors. These variants can increase the specificity of library searches. We argue that these CSA spectra should be catalogued in a mass spectral library and in a community driven MS database such as GNPS^10^. A CSA mass spectral library is a collection of unique composite spectra variants of different chemical compounds which can be created using MS1 only data for annotated peaks and authentic standards. To create CSA libraries, raw LC/HRMS data for known annotations or authentic standards are needed, which are readily available for over 2000 publicly available metabolomics datasets in EBI Metabolights^24^, GNPS Massive^25^ and Metabolomics Workbench^26^ repositories.

Our work is inspired by RAMClust^8^. However, we have added new features 1) LOESS based smoothing for extracted ion chromatograms 2) the input is IDSL.IPA MS1 peaklists which only have pre-filtered ^12^ C peaks 3) spectra annotation using both entropy and cosine similarity scores 4) filtering of CSA spectra using m/z range, fragment count, inter-and intra-sample correlations 5) all functions are coded in R with minimal dependencies on other external R packages 6) a user-friendly excel sheet for parameter selection and reproducible data processing and 7) multi-threaded computation for Linux and Windows OS.

The IDSL.CSA workflows incorporate several essential and novel features for processing untargeted metabolomics datasets. It encapsulates a full workflow of steps including peak detection^19^, alignment, DDA/DIA deconvolution, library generation, spectra search and annotation ranking, into a single line of R command that need all parameters in an input Microsoft Excel file. The workflow applies a critical step of LOESS smoothing followed by the cubic spline smoothing method to minimize chromatogram jaggedness while computing correlation among ion intensities. CSA variants and consensus CSA spectra both were created for a compound to enable cross-instrument searches. It also interprets the CSA spectrum by identifying ion species of commonly observed ESI adducts. The optimized pre-filtering methods using precursor, spectra markers, spectra entropy enabled faster searching of larger libraries with millions of spectra. A key unique strength of our approach is to identify recurring CSA spectra across multiple samples which is helpful in generating high confidence CSA libraries. It also standardizes publicly available mass spectra data with inconsistent fields to a standard storage format in R, making IDSL.CSA and IDSL.FSA packages fully compatible with existing public MS/MS libraries and the NIST MS/MS database. With these features, IDSL.CSA workflow can help in generating high-quality data matrices from untargeted metabolomics datasets. IDSL.CSA is a useful addition to the growing pool of software to improve the annotation of MS1 data in untargeted metabolomics studies.

It is a major challenge in metabolomics that 2/3 of untargeted studies in the Metabolomics Workbench data repository (https://www.metabolomicsworkbench.org) have only unannotated peaks. To overcome this hurdle to some extent, IDSL.CSA workflows may be able to annotate a substantial number of peaks in these studies using available reference MS/MS and newly created CSA libraries. With the emergence of these data repositories, it is possible to process a large number of untargeted datasets to capture different variants of CSA spectra that may originate due to variations in experimental conditions. By utilizing MS1 data in an exhaustive way, our workflow can minimize the underutilization of untargeted metabolomics datasets in studying basic metabolic processes and biomonitoring of environmental chemical exposure.

## Author’s contribution

SFB and DKB planned the study, prepared the results and drafted the manuscript. SFB and DKB coded the IDSL.CSA package. YK provided test LC/HRMS data for authentic standards of metabolites. All authors have reviewed the manuscript content.

## Supporting information

Supporting spreadsheet tables, data sources, workflow and computational method details and additional benchmarks.

