## Supplementary Material for "IDSL.CSA: Composite Spectra Analysis for Chemical Annotation of Untargeted Metabolomics Datasets"

**Table of Content:**

**Table S.1.** LC/HRMS datasets used in this work

**S.1. Comparison between peaklist and aligned table CSA analysis**

**Figure S.1.1.** Comparison between CSA aligned EICs and pseudo-spectra of Hippuric acid and Phenylalanyl-Hydroxyproline that were deconvoluted using two built-in CSA approaches.

**Figure S.1.2.** Pairwise Pearson’s correlation table of the heights of Hippuric acid (*m/z* = 180.066) and Phenylalanyl-Hydroxyproline (*m/z* = 261.124) ions on the aligned peak height table. (^13^C isotopologue peaks were excluded from this table, and coefficients ≤ 0.5 were shown by 0.

**Figure S.1.** Mass spectral similarity search results for the CSA spectra of file pool_P1_A1-A12.mzML from MSV000088661 (IROA standards) study.

**Figure S.2.** Mass spectral similarity search results for the DDA spectra of file pool_P1_A1-A12.mzML from MSV000088661 (IROA standards) study.

**Figure S.3.** Comparison between CSA and DDA spectra integration methods for kynurenine (RT = 4.167) from MSV000088661 (IROA standards) study.

**Figure S.4.** Comparison between two composite spectra variants of kynurenine in two standard samples from MSV000088661 (IROA standards) study.

**S.2. Benchmarking the spectra search filtering with IDSL.FSA**

**Figure S.2.1.** Schematic of spectra marker selection procedure for tryptophan in an authentic standard mixture from MSV000088661 study.

**Figure S.2.2.** Distribution of absolute spectral entropy differences (ΔS) across 272 authentic standard compounds in a DDA analysis from the MSV000088661 study.

**Table S.2.1.** Evaluation of pre-filtering steps for a library 1.8 million spectra

**Table S.2**. Parameters for CSA spectra extraction

**Table S.3.** Parameters for DIA spectra extraction

**Table S.4.** Parameters for DDA spectra extraction

**Table S.5.** Parameters for spectral searches

| **Table S.1.** LC/HRMS datasets used in this work | | | | | |
| --- | --- | --- | --- | --- | --- |
| **Sample Type** | **Repository** | **Accession ID** | **Analysis mode** | **Mass analyzer** | **Number of samples** |
| Neat standard compounds | EBI metabolights | MTBLS1040 | HILIC – ESI – POS | Agilent 6550 Q-TOF | 978 |
| IROA standards | GNPS | MSV000088661 | RP – ESI – POS | Thermo Fusion Orbitrap | 54 |
| Cardiovascular disease | MetabolomicsWorkbench | ST002044 | RP – ESI – POS | Thermo Fusion Orbitrap | 286 |
| Human microbiome | MetabolomicsWorkbench | ST000923 | HILIC/RP – ESI – POS/NEG | Thermo Q Exactive Orbitrap | 600 |
| Gut microbiome | MetabolomicsWorkbench | ST001000 | HILIC/RP – ESI – POS/NEG | Thermo Q Exactive Orbitrap | 226 |

**S.1. Comparison between peaklist and aligned table CSA analysis**

Hippuric acid and Phenylalanyl-Hydroxyproline elute simultaneously in the MTBLS2295 study acquired using a SeQuant ZIC-HILIC column. When only individual peaklists are used, these two compounds are grouped in the same composite spectra block. However, when peaks that already correlated at the height aligned tables level are used for the composite spectra analysis, these two compounds are separated with no interferences shown in Figure S.1.1.

| 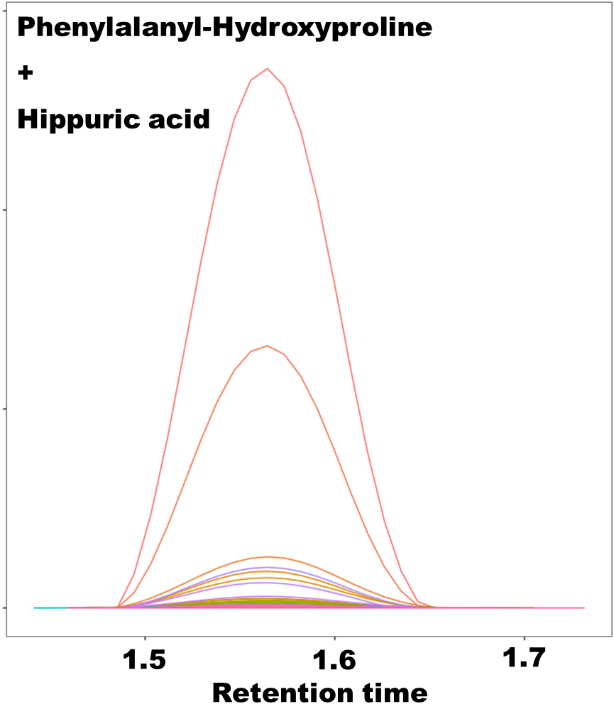 | 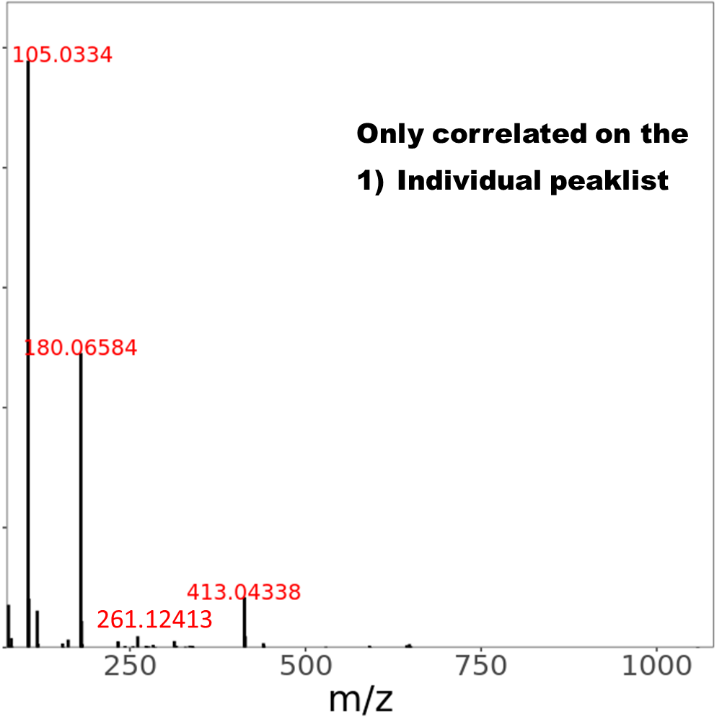 |
| --- | --- |
| 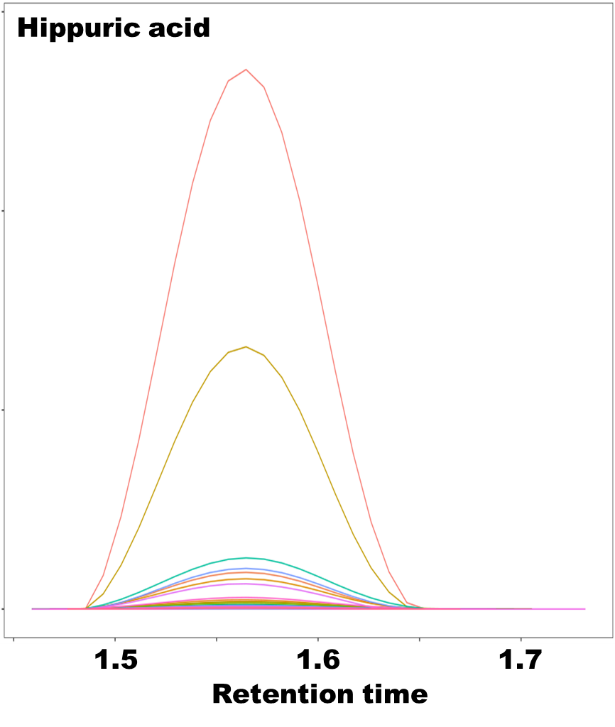 | 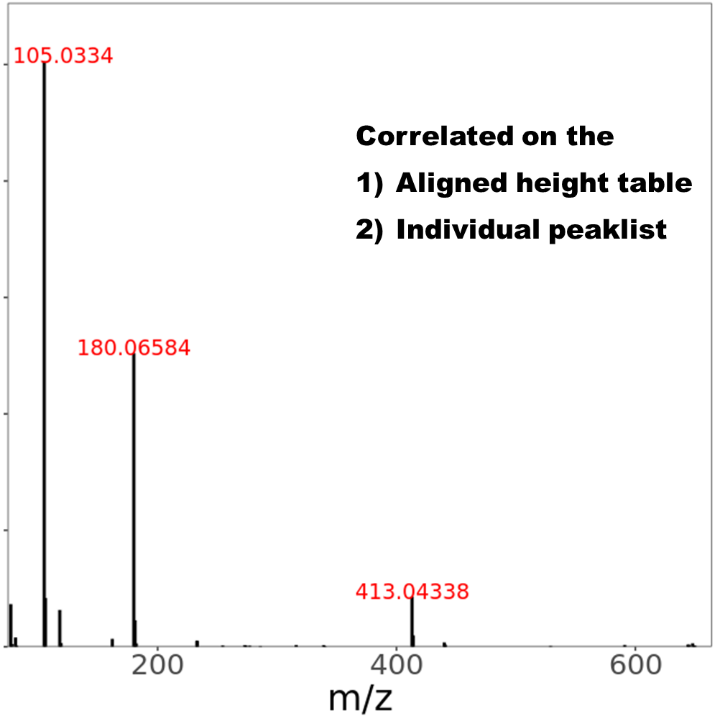 |
| 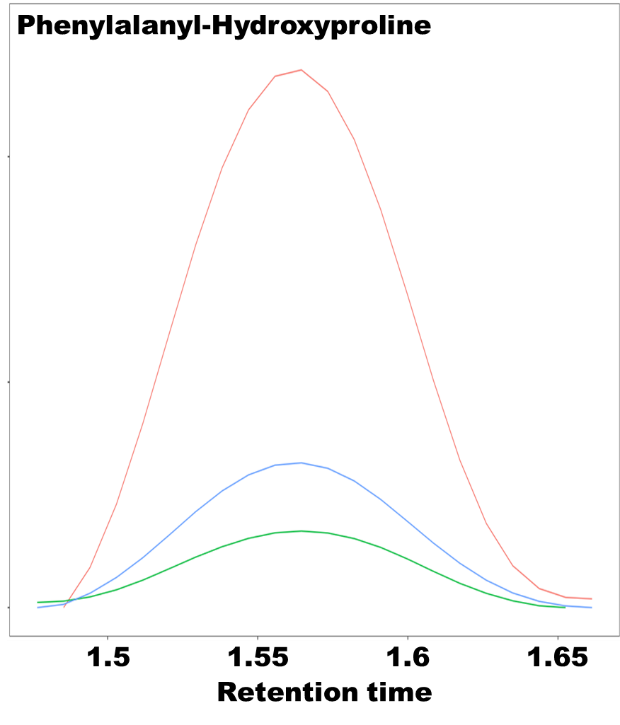 | 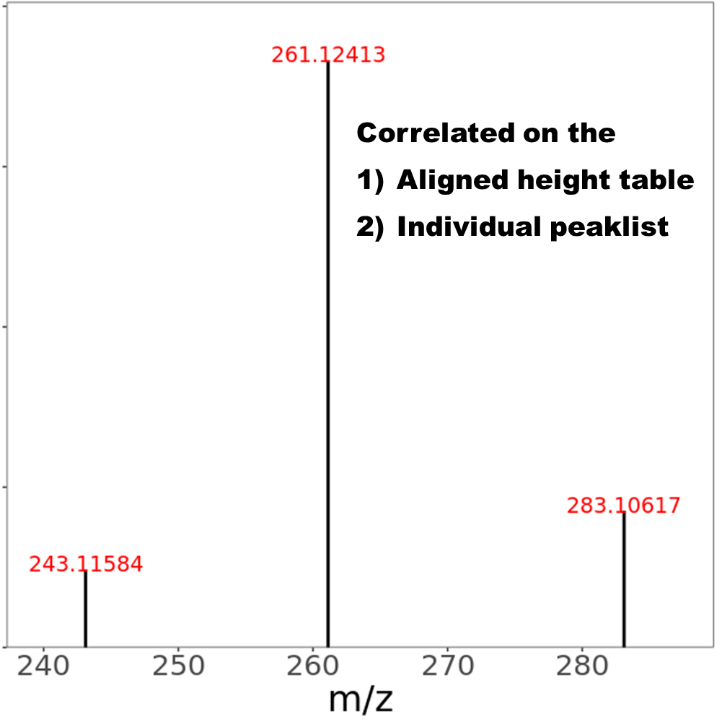 |
| **Figure S.1.1.** Comparison between CSA aligned EICs and pseudo-spectra of Hippuric acid and Phenylalanyl-Hydroxyproline that were deconvoluted using two built-in CSA approaches. | |

The composite spectra approach of the only peaklist corrolation incorrectly merged the Phenylalanyl-Hydroxyproline (261.12413) peaks with the Hippuric acid peaks due to strong co-elution. However, when aligned table corrolation was used, Hippuric acid and Phenylalanyl-Hydroxyproline were well seperated shown in Figure S.1.1. Corroloation of these peaks on the aligned table were investigated and shown on Figure S.1.2, and shown the 261.124 and 180.066 do not have any corrolation on the aligned height table.

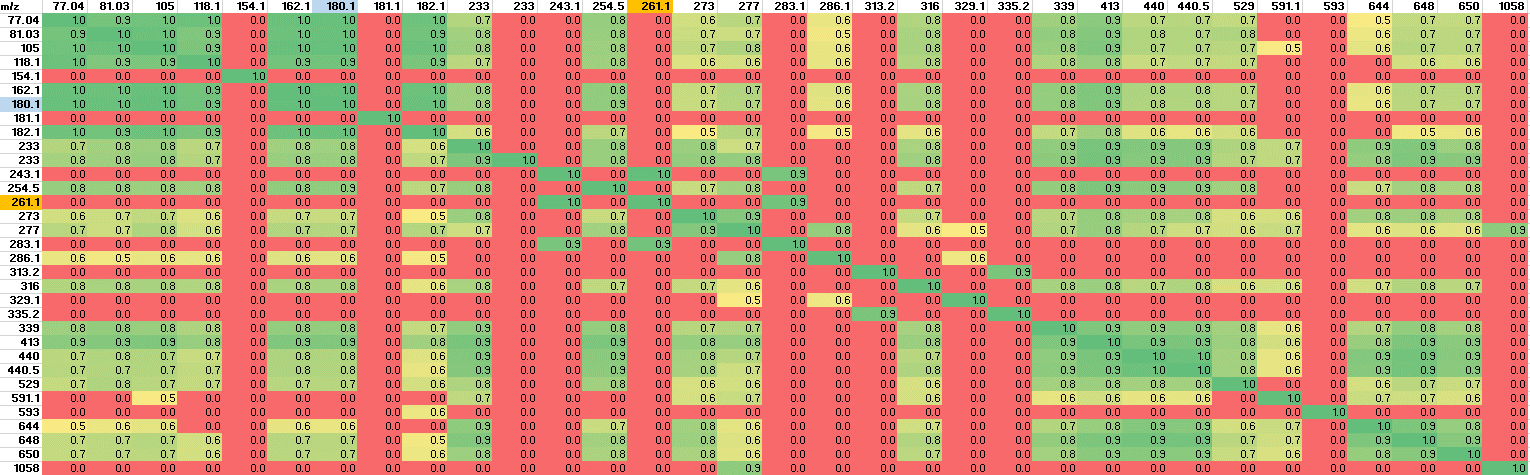

**Figure S.1.2.** Pairwise Pearson’s correlation table of the heights of Hippuric acid (m/z = 180.066) and Phenylalanyl-Hydroxyproline (m/z = 261.124) ions on the aligned peak height table. (^13^C isotopologue peaks were excluded from this table, and coefficients ≤ 0.5 were shown by 0.

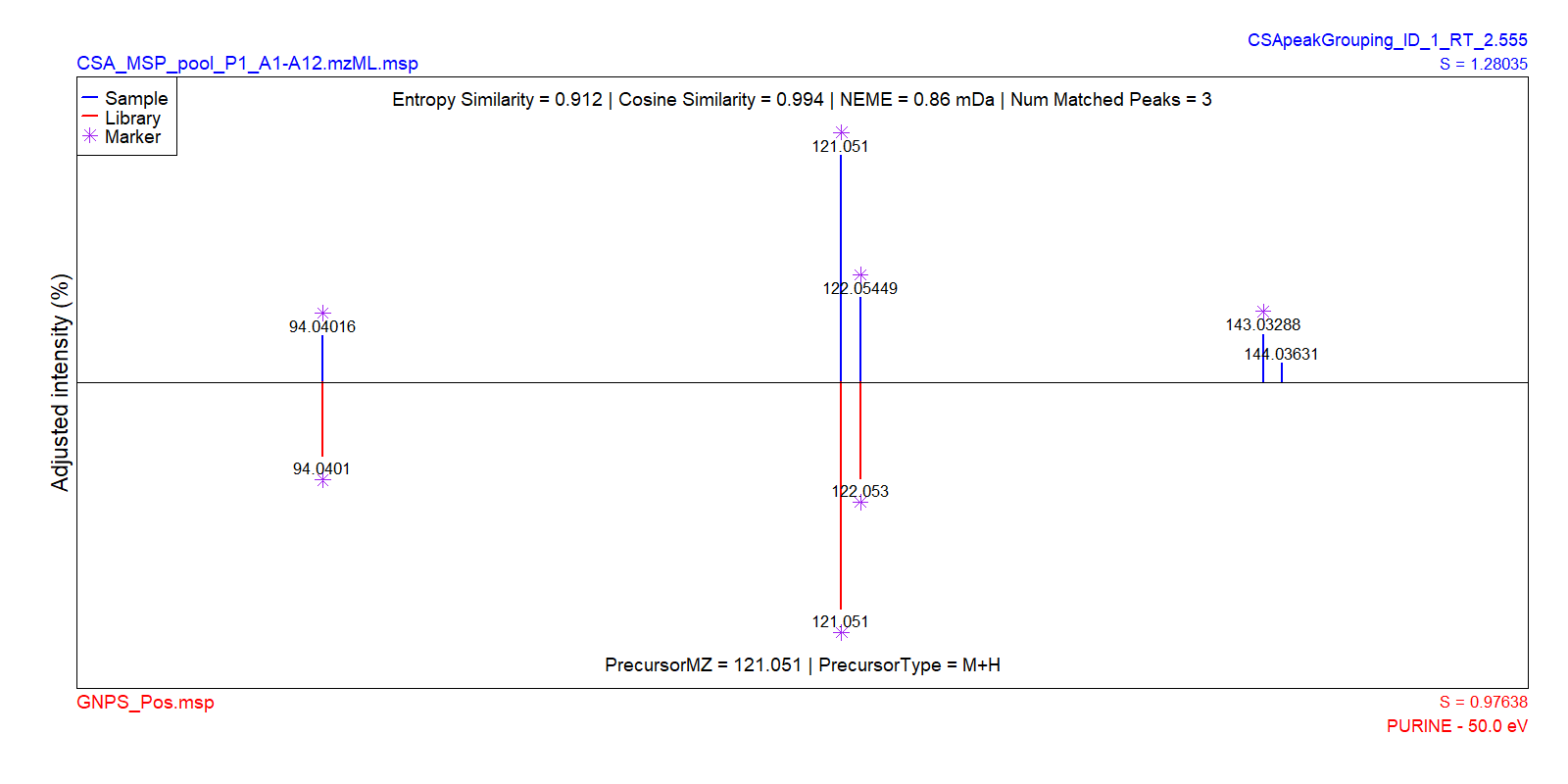

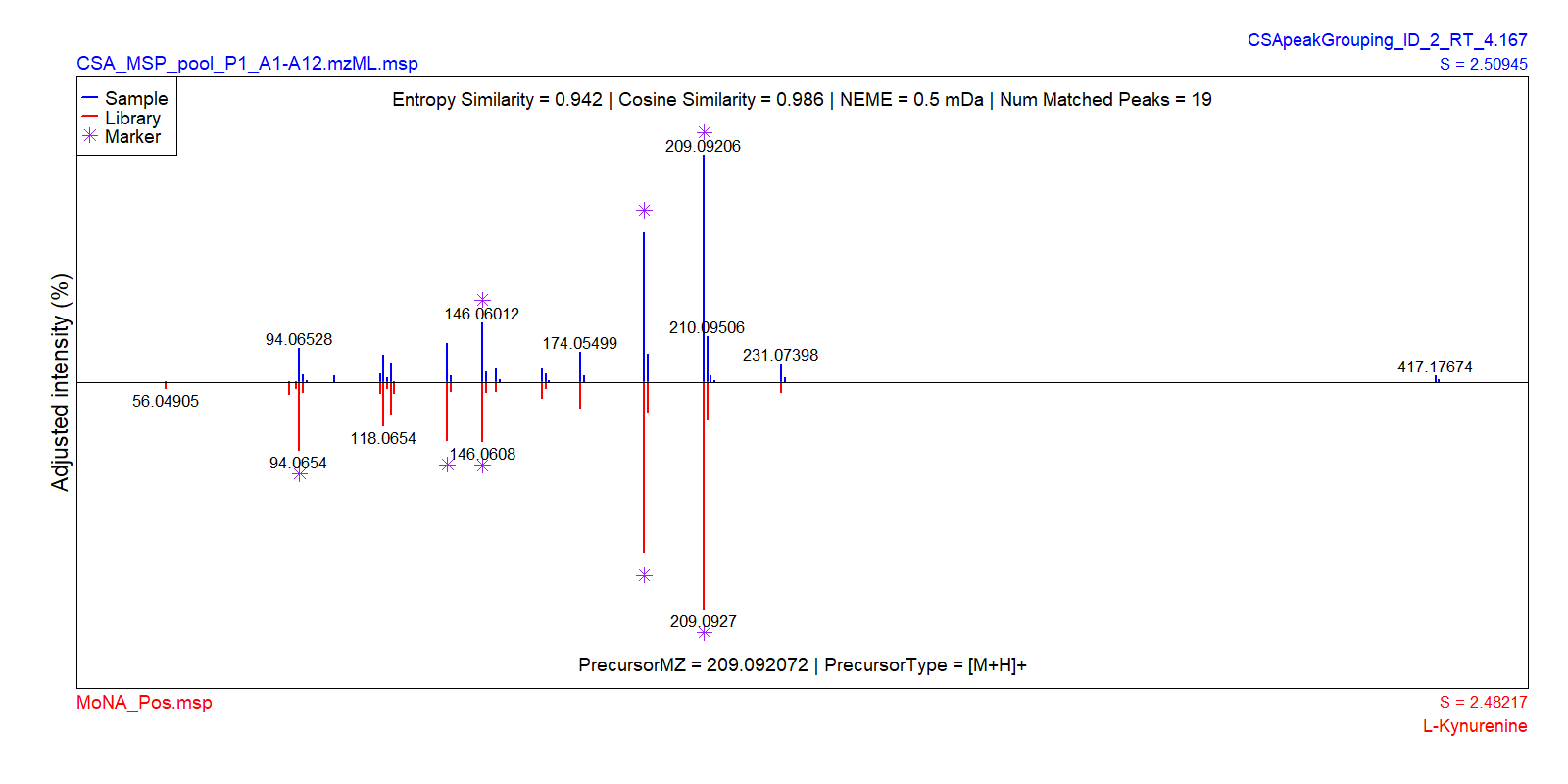

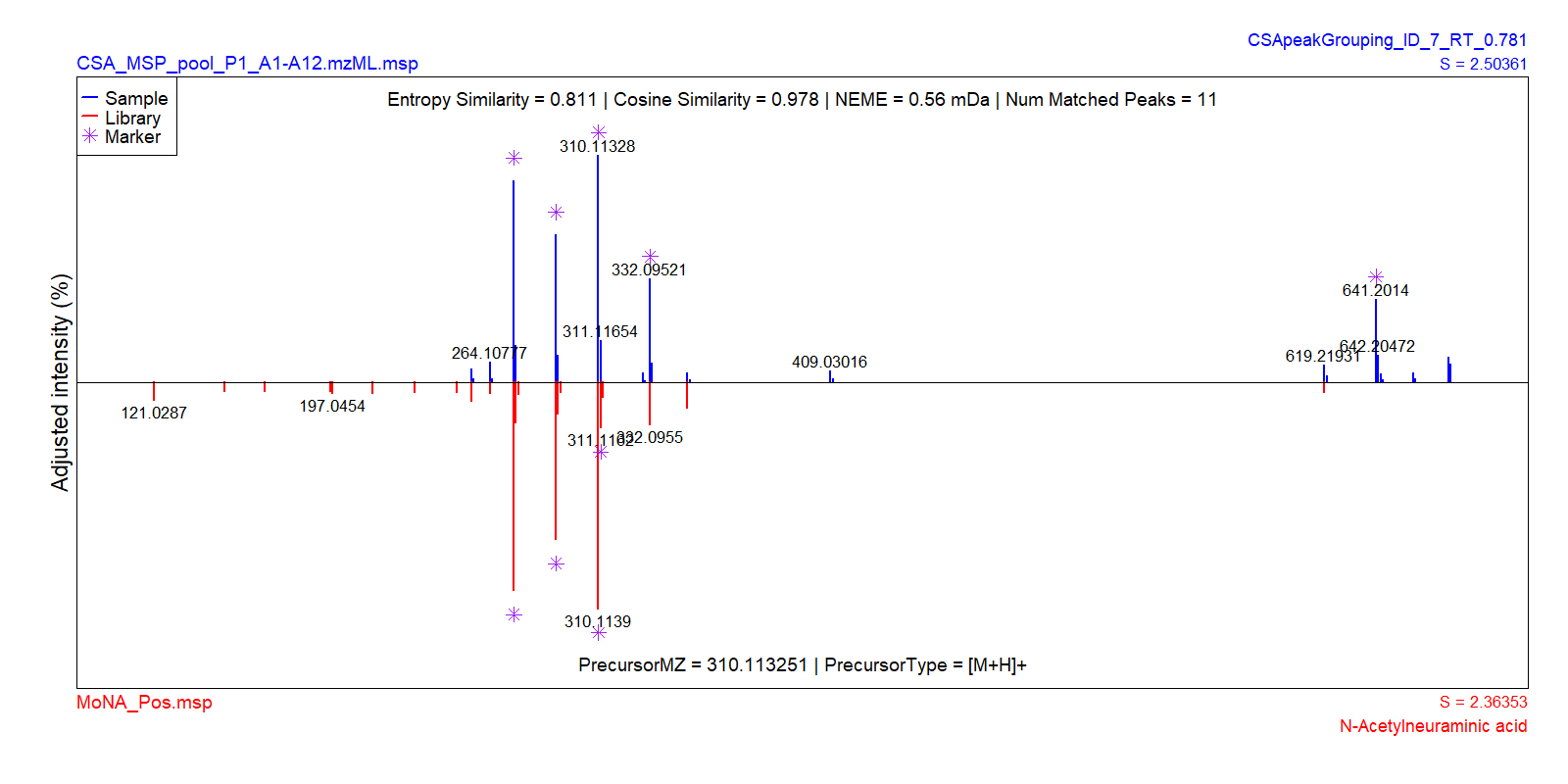

**Figure S.1.** Mass spectral similarity search results for the CSA spectra of file pool_P1_A1-A12.mzML from MSV000088661 (IROA standards) study. (Noise removal = 0.5%)

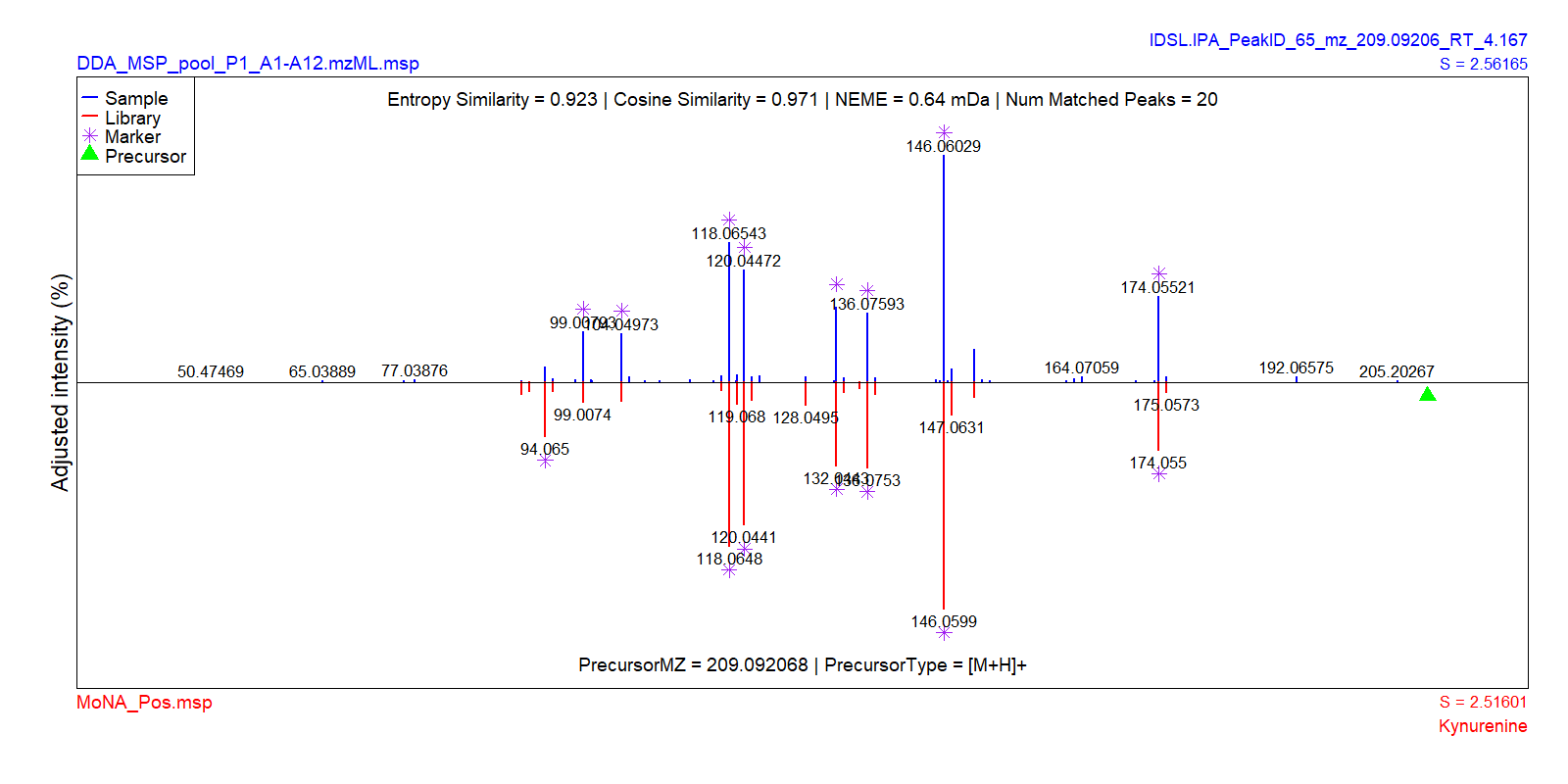

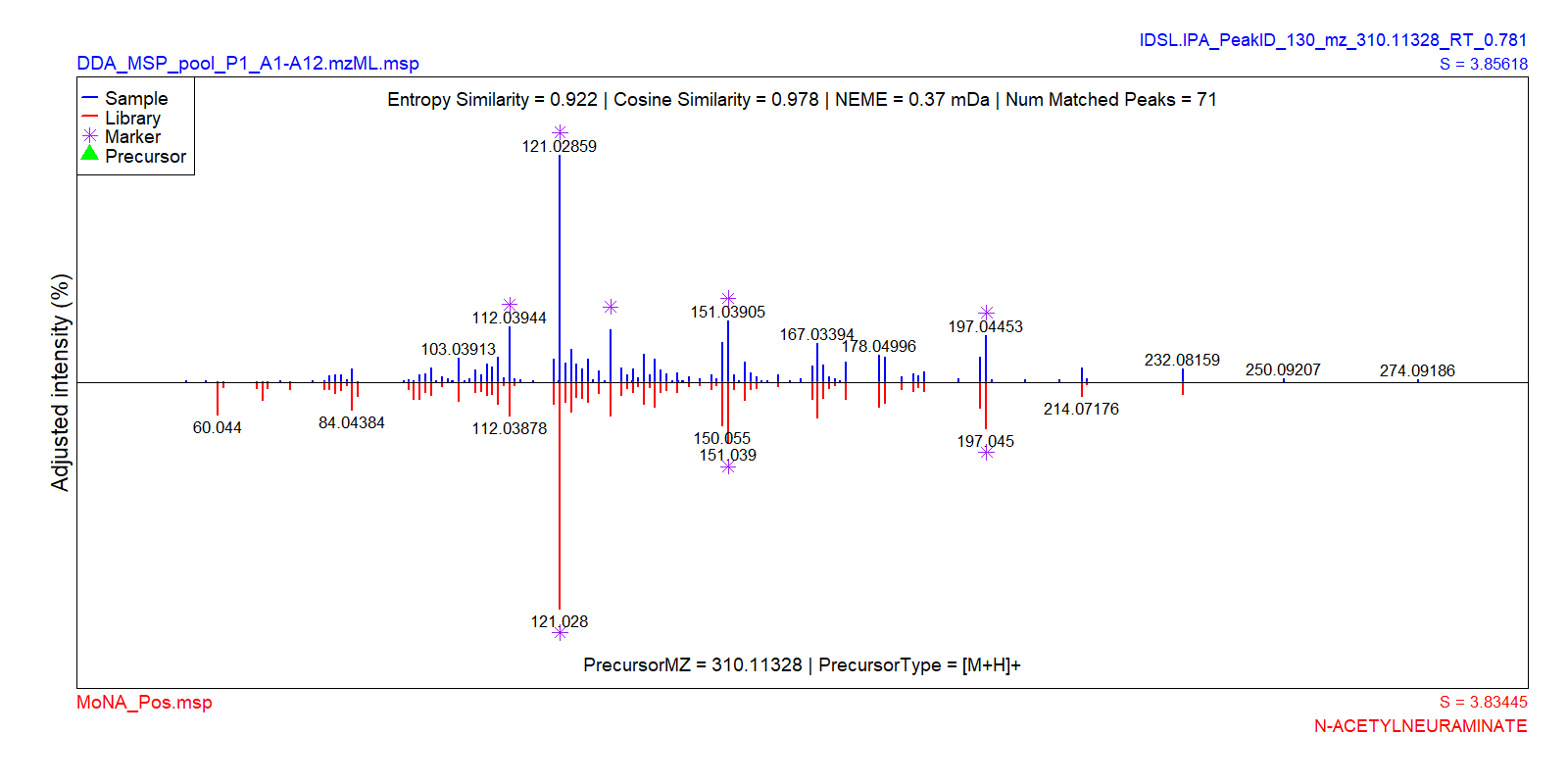

**Figure S.2.** Mass spectral similarity search results for the DDA spectra of file pool_P1_A1-A12.mzML from MSV000088661 (IROA standards) study.

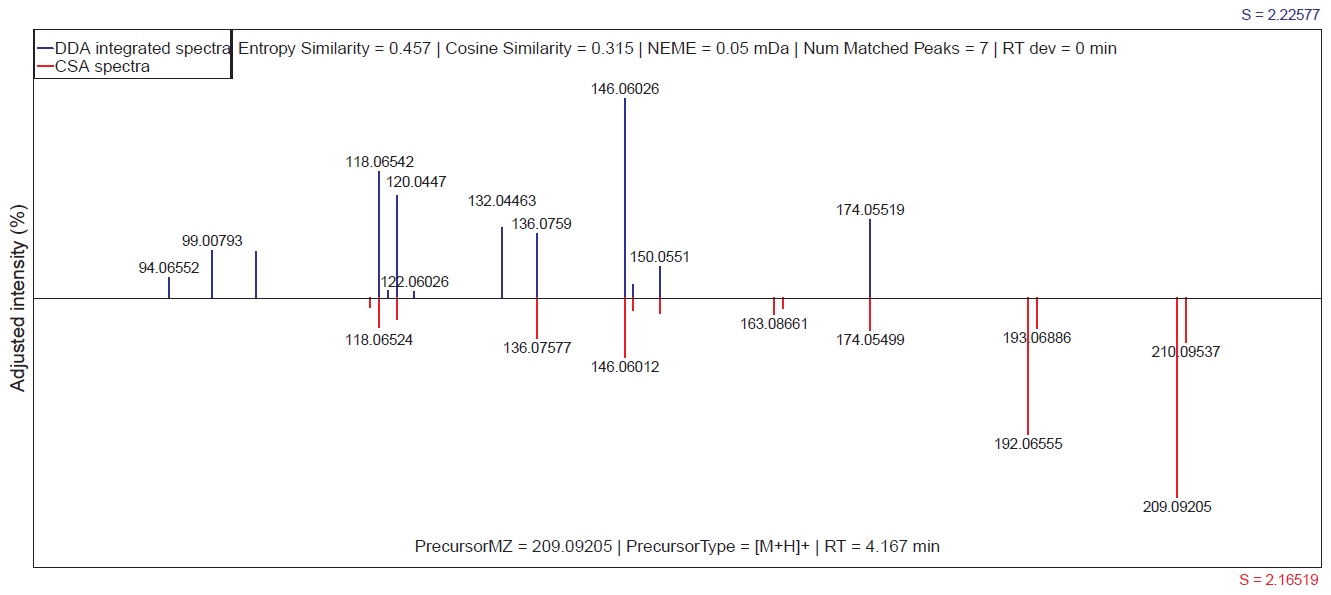

**Figure S.3.** Comparison between CSA and DDA spectra integration methods for kynurenine (RT = 4.167) from MSV000088661 (IROA standards) study.

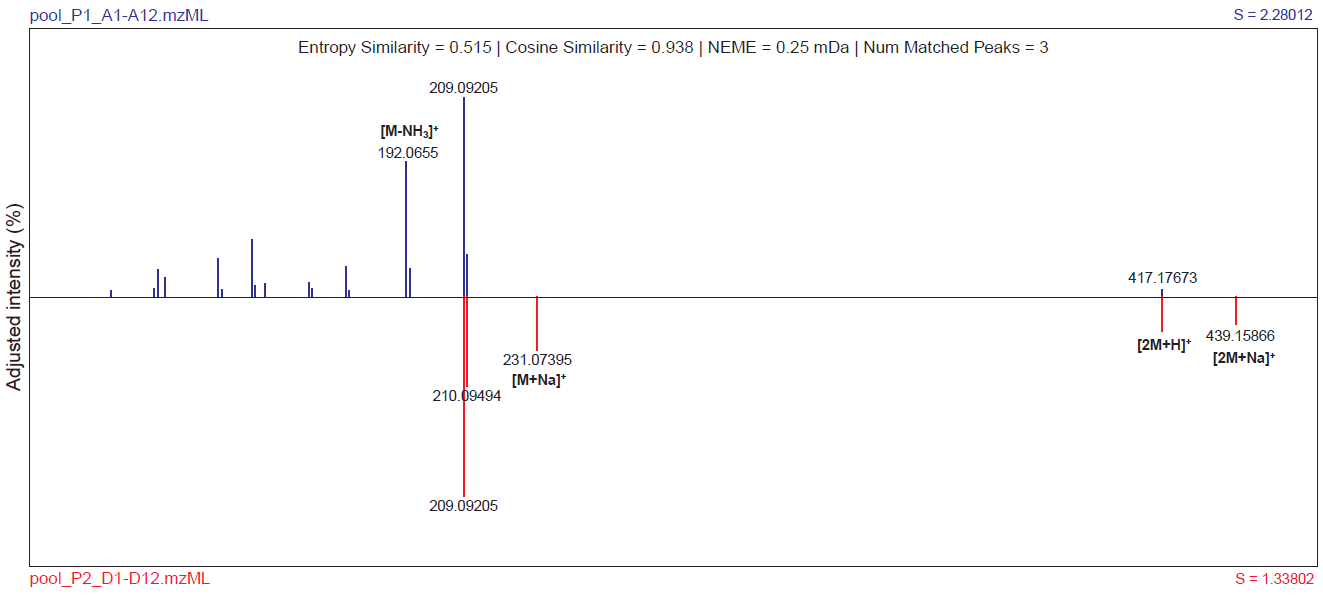

**Figure S.4.** Comparison between two composite spectra variants of kynurenine in two standard samples from MSV000088661 (IROA standards) study.

**S.2. Benchmarking the spectra search filtering with IDSL.FSA**

IDSL.FSA includes five optional pre-filtering steps to narrow down lists of candidate reference fragmentation patterns to accelerate the matching procedure. IDSL.FSA required 1.5 hours to correctly match one MSP block of tryptophan on a combined library of GNPS, MoNA, and NIST 20 containing 1.8 million spectra without any pre-filtering steps on a single-thread processor.

1) **Spectra markers**

Spectra markers are characteristic ions above a baseline in the reference and sample fragmentation patterns. References fragmentation patterns with a good percentage of matched nominal or rounded masses are selected for the next step of annotation. Figure S.2.1 presents schematic of ion marker selection process for tryptophan fragmentation pattern.

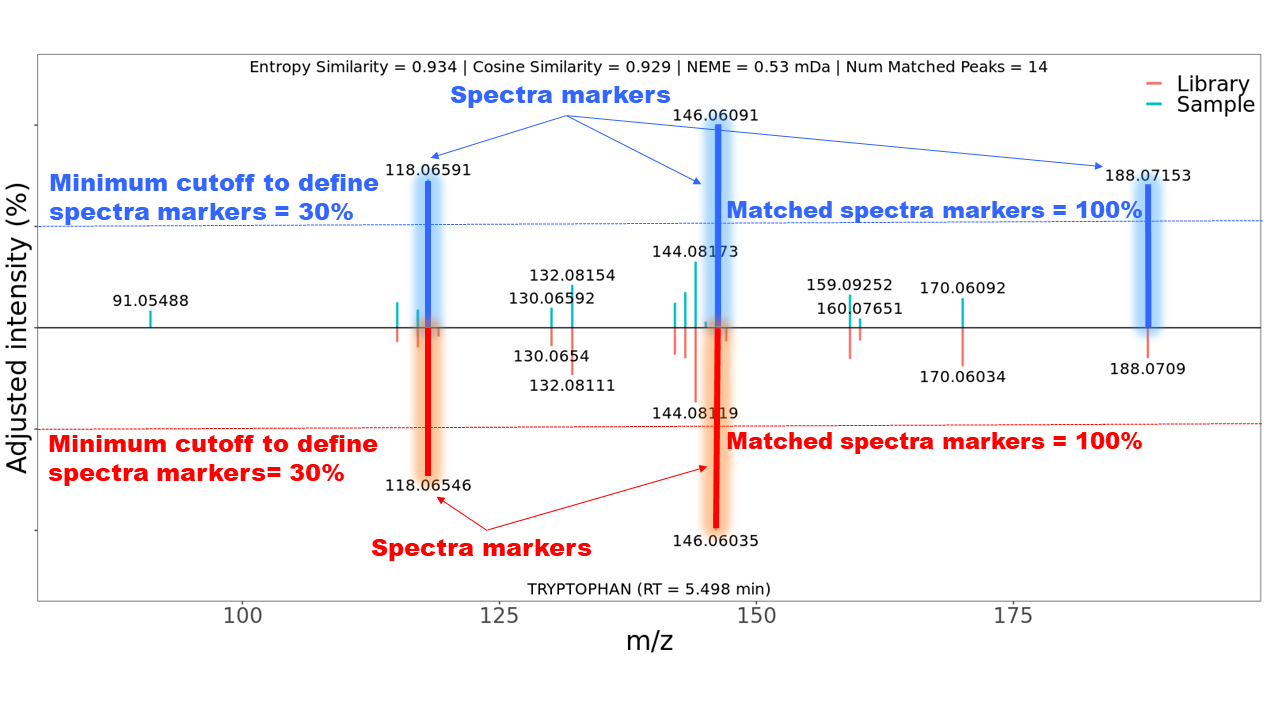

**Figure S.2.1.** Schematic of spectra marker selection procedure for tryptophan in an authentic standard mixture from MSV000088661 study.

Only pre-filtering step using spectra markers was able to reduce number of reference fragmentation patterns for tryptophan to 2213 hits with baseline cutoff 20% and percentage of matched spectra of 50% after applying rounding digit of 1 only on spectra markers.

2) **Absolute spectral entropy differences**

Prior to matching fragmentation spectra, IDSL.FSA calculates spectral entropy values. Absolute differences of these raw values can also implemented in the pre-filtering step. Figure S.2.2 presents distribution of absolute spectral entropy differences (ΔS) across 272 compounds in a DDA analysis from MSV000088661 study.

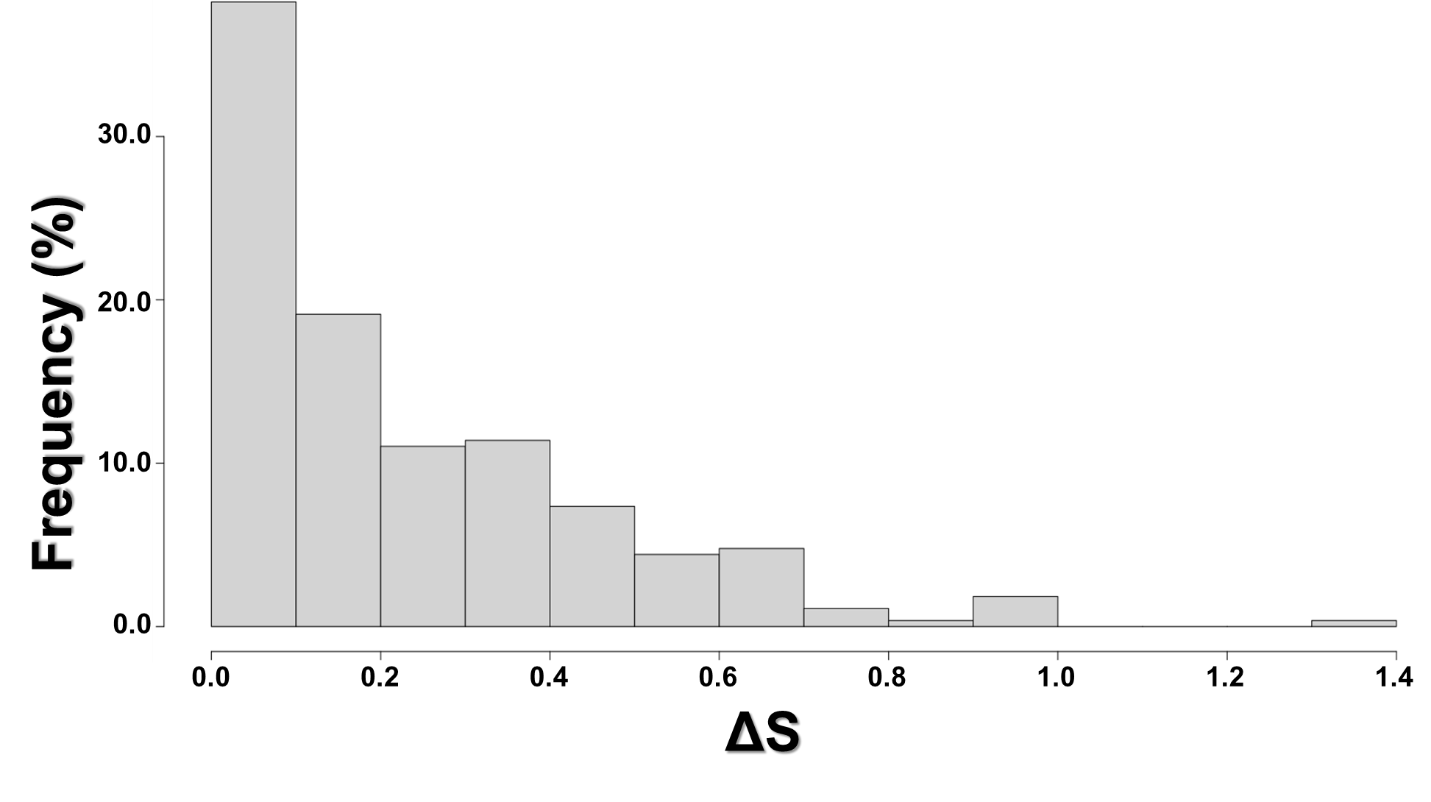

**Figure S.2.2.** Distribution of absolute spectral entropy differences (ΔS) across 272 authentic standard compounds in a DDA analysis from the MSV000088661 study.

Only pre-filtering step using absolute spectral entropy differences (ΔS) was able to reduce number of reference fragmentation patterns for tryptophan to 338264 hits with ΔS ≤ 1.4.

3) **Precursor m/z match**

Precursor m/z match is optional for the IDSL.FSA workflow. This option allows matching fragmentation patterns that do not have reliable precursor values. Only pre-filtering step using precursor matching with mass window ≤ 0.01 *Da* was able to reduce number of reference fragmentation patterns for tryptophan to 415 hits.

4) **Retention time match**

When retention time values were acquired from in-house instruments, retention time pre-filtering can be implemented for IDSL.FSA to further narrow down the list of candidate reference fragmentation patterns. This pre-filtering step was not benchmarked in this work since retention time data from this public reference library were not applicable to this work.

5) **Precursor adduct type**

Precursor adduct types may be implemented to only perform annotation on a selected list of adduct types. IDSL.FSA is able to curate various adduct terminologies into a standard format to optimally parse fragmentation libraries.

The minimum required time to reach pre-processing stage for this library was 51.98632 seconds. This pre-processing step is executed once for each run and do not need to be repeated to annotate other MSP blocks. A combination of these pre-filtering steps resulted with 191 hits to match tryptophan on this library which only required 1.8 seconds on a single-thread processor indicating scalability of IDSL.FSA for population-size studies. The IDSL.FSA pre-filtering speed improvements were benchmarked in Table S.2.1.

| **Table S.2.1.** Evaluation of pre-filtering steps for a library 1.8 million spectra | | |
| --- | --- | --- |
| **Pre-filtering step** | **Number of candidate reference fragmentation patterns** | **Required processing time** |
| **Spectra markers** | 2213 | 3.8451 seconds |
| **Absolute spectral entropy differences** | 338264 | 4.751767 minutes |
| **Precursor m/z match** | 415 | 35.50688 seconds |
| **Combined pre-filtering steps** | 191 | 1.84055 seconds |

**Table S.2.** Parameters for CSA spectra extraction

| **Analysis step** | **Parameter ID** | **Parameter description** | **User provided input** |
| --- | --- | --- | --- |
| **Global CSA (required)** | CSA0001 | Process individual IDSL.IPA peaklist | YES |
|  | CSA0002 | Create a library of unique spectra variants | YES |
|  | CSA0003 | Aggregate CSA spectra on the aligned table | YES |
|  | CSA0004 | Number of parallel threads (only on "Sample Mode" levels) | 1 |
| **Data import and export (required)** | CSA0005 | HRMS data location | /path/to/folder/ |
|  | CSA0006 | Address of the reference standard metadata file to generate a list of targeted CSA blocks | NA |
|  | CSA0007 | List of HRMS files (.mzXML/.mzML/.CDF) | All |
|  | CSA0008 | Address of the `peaklists` directory generated by the IDSL.IPA workflow. | /peaklists |
|  | CSA0009 | **Optional:** Address of the `peak_alignment` directory generated by the IDSL.IPA workflow. This folder should be generated through the peak alignment modules of the IDSL.IPA pipeline (***PARAM0002 in the IDSL.IPA parameter spreadsheet***). | /peak_alignment |
|  | CSA0010 | Generate aligned extracted ion chromatograms (EICs) figures for deconvoluted ions | NO |
|  | CSA0011 | Output location (*.msp* files and EICs) | /path/to/folder/ |
| **Individual CSA *.msp* generation** | CSA0012 | Select **peakList** or **alignedTable** to indicate the source of reoccurring clusters of peaks. **peakList** implies that the inter-sample correlation among peaks is used to cluster related peaks. **alignedTable** means to cluster related peaks that also correlate on the aligned peak height table. | peakList |
|  | CSA0013 | Retention time window (min) to find reoccurring peaks in the IDSL.IPA peaklists | 0.01 |
|  | CSA0014 | A minimum baseline S/N threshold for the most abundant m/z peaks from the IDSL.IPA peaklist | 5 |
|  | CSA0015 | Smoothing window (number of scans for smoothing) for MS1 channel | 25 |
|  | CSA0016 | Mass accuracy (Da) | 0.005 |
|  | CSA0017 | Number of extra scans on the both sides of detected peak boundaries to produce EICs more accurately | 250 |
|  | CSA0018 | Number of points to smooth individual chromatographic peaks using cubic spline method to increase number of data points for the Pearson's correlation | 2000 |
|  | CSA0019 | Percentage of top height of the chromatographic peaks to measure peak correlation similarities (%) | 100 |
|  | CSA0020 | Minimum distance (Da) between lowest and highest m/z to prevent clustering only isotopic envelopes | 0 |
|  | CSA0021 | Minimum number of 12C ions in a CSA cluster | 2 |
|  | CSA0022 | Pearson’s correlation coefficient threshold | 0.99 |
| **Creat unique spectra by measuring pairwise similarity across entire samples** | CSA0023 | Aggregation meta-variable (only when **CSA0006** is provided) | Name |
|  | CSA0024 | Export classified spectra plots for individual unique spectra | NO |
|  | CSA0025 | Mass window (Da) to match reference spreadsheet compounds | 0.005 |
|  | CSA0026 | Retention time window (min) to find CSA spectra variants or match reference spreadsheet compounds | 0.05 |
|  | CSA0027 | Minimum frequency (%) of detection for CSA spectra across all *.msp*files for untargeted CSA spectra | 10 |
|  | CSA0028 | Weighted spectra to measure entropy similarity score. Weighted functions are used to boost the intensity of low abundant peaks described by Li *et. al.,* **2021**, *Nature Methods* | TRUE |
|  | CSA0029 | Minimum spectral entropy similarity score to find similar CSA spectra | 0.9 |
| **CSA aggregation on the aligned table** | CSA0030 | Retention time window (min) to group reoccurring peaks | 0.05 |
|  | CSA0031 | Minimum frequency (%) of detection for CSA peaks across the aligned peak table | 5 |
|  | CSA0032 | Minimum number of reoccurring fragments in each cluster | 3 |
|  | CSA0033 | Minimum Tanimoto coefficient to cluster reoccurring peaks | 0.1 |
|  | CSA0034 | Minimum Tanimoto coefficient to generate integrated and the most abundant individual CSA spectra .msp files. An integrated spectra contains a median of intensities for reoccurring peaks across samples in which peaks were detected. An abundant spectra is the CSA spectra from the data file in which the most abundant fragment ion occured. This parameter controls number of CSA spectra variants. | 0.5 |
|  | CSA0035 | Export aligned CSA spectra plots | NO |
|  | CSA0036 | Mass accuracy (Da) to create 1:1 mixing spectra for spectral entropy similarity | 0.005 |
|  | CSA0037 | Weighted spectra to measure entropy similarity score. Weighted functions are used to boost the intensity of low abundant peaks described by Li *et. al.,* **2021**, *Nature Methods* | TRUE |
|  | CSA0038 | Minimum spectral entropy similarity score for mass spectral similarity network creation | 0.6 |
|  | CSA0039 | Address of an **FSDB** to annotate *.msp* network of spectra using default values | NA |

**Table S.3.** Parameters for DIA spectra extraction

| **Analysis step** | **Parameter ID** | **Parameter description** | **User provided input** |
| --- | --- | --- | --- |
| **Global DIA (required)** | DIA0001 | Process individual IDSL.IPA peaklist | YES |
|  | DIA0002 | Create a library of unique spectra variants | YES |
|  | DIA0003 | Number of parallel threads | 1 |
| **Data import and export (required)** | DIA0004 | HRMS data location | /path/to/folder/ |
|  | DIA0005 | Address of the reference standard metadata file to generate a list of targeted DIA blocks | NA |
|  | DIA0006 | List of HRMS files (.mzXML/.mzML/.CDF) | All |
|  | DIA0007 | Address of the `peaklists` directory generated by the IDSL.IPA workflow. | /peaklists |
|  | DIA0008 | **Optional:** Address of the `peak_alignment` directory generated by the IDSL.IPA workflow. This folder should be generated through the peak alignment modules of the IDSL.IPA pipeline (***PARAM0002 in the IDSL.IPA parameter spreadsheet***). | /peak_alignment |
|  | DIA0009 | Index number(s) of selected IDSL.IPA peaks | All |
|  | DIA0010 | Generate aligned extracted ion chromatograms (EICs) figures for deconvoluted ions | NO |
|  | DIA0011 | Output location (*.msp* files and aligned EICs figures) | /path/to/folder/ |
| **DIA *.msp* generation** | DIA0012 | Parallelization mode | Sample Mode |
|  | DIA0014 | Intensity height threshold for chromatographic peaks | 1.00E+03 |
|  | DIA0015 | Smoothing window (number of scans for smoothing) for MS1 channel | 15 |
|  | DIA0016 | Smoothing window (number of scans for smoothing) for MS2 channel | 12 |
|  | DIA0017 | Mass accuracy (Da) | 0.01 |
|  | DIA0018 | Number of extra scans on the both sides of detected peak boundaries to produce EICs more accurately | 50 |
|  | DIA0019 | Number of points to smooth individual chromatographic peaks using cubic spline method to increase number of data points for the Pearson's correlation | 100 |
|  | DIA0020 | Percentage of top height of the chromatographic peaks to measure peak correlation similarities (%) | 90 |
|  | DIA0021 | Pearson’s correlation coefficient threshold | 0.95 |
| **Creat unique spectra by measuring pairwise similarity across entire samples** | DIA0022 | Aggregation meta-variable (only when **DIA0005** is provided) | Name |
|  | DIA0023 | Export classified spectra plots for individual unique spectra | NO |
|  | DIA0024 | Mass window (Da) to match reference spreadsheet compounds | 0.01 |
|  | DIA0025 | Retention time window (min) to find DIA spectra variants or match reference spreadsheet compounds | 0.05 |
|  | DIA0026 | Minimum frequency (%) of detection for DIA spectra across all *.msp* files for untargeted DIA spectra | 10 |
|  | DIA0027 | Weighted spectra to measure entropy similarity score. Weighted functions are used to boost the intensity of low abundant peaks described by Li *et. al.,* **2021**, *Nature Methods* | TRUE |
|  | DIA0028 | Minimum spectral entropy similarity score to find similar DIA spectra | 0.9 |

**Table S.4.** Parameters for DDA spectra extraction

| **Analysis step** | **Parameter ID** | **Parameter description** | **User provided input** |
| --- | --- | --- | --- |
| **Global DDA (required)** | DDA0001 | Process individual IDSL.IPA peaklist | YES |
|  | DDA0002 | Create a library of unique spectra variants | YES |
|  | DDA0003 | Number of parallel threads | 1 |
| **Data import and export (required)** | DDA0004 | HRMS data location | /path/to/folder/ |
|  | DDA0005 | Address of the reference standard metadata file to generate a list of targeted DDA blocks | NA |
|  | DDA0006 | List of HRMS files (.mzXML/.mzML/.CDF) | All |
|  | DDA0007 | Address of the `peaklists` directory generated by the IDSL.IPA workflow. | /peaklists |
|  | DDA0008 | **Optional:** Address of the `peak_alignment` directory generated by the IDSL.IPA workflow. This folder should be generated through the peak alignment modules of the IDSL.IPA pipeline (***PARAM0002 in the IDSL.IPA parameter spreadsheet***). | /peak_alignment |
|  | DDA0009 | Index number(s) of selected IDSL.IPA peaks | All |
|  | DDA0010 | Generate DDA spectra figures | NO |
|  | DDA0011 | Output location (*.msp* files and spectra figures) | /path/to/folder/ |
| **DDA *.msp* generation** | DDA0012 | Parallelization mode | Sample Mode |
|  | DDA0013 | Mass accuracy (Da) to find precursor masses in the IDSL.IPA peaklist | 0.01 |
|  | DDA0014 | ***Integrate***, ***filter***, or select ***the most abundant*** of DDA spectra (MS2) of the same precursor across MS1 chromatographic peaks (mass spectra). **IonFiltering** can only be applied when DDA scan count ≥ 3; however, the "**MostIntenseDDAspectra"** method is still used when DDA scan counts ≤ 2. | IonFiltering |
|  | DDA0015 | Mass accuracy (Da) for "**IonFiltering**" or "**DDAspectraIntegration**" | 0.01 |
|  | DDA0016 | Minimum percentage of detected scans to calculate Pearson's correlation coefficient. A minimum number of 3 detected DDA scans automatically is applied across MS1 chromatographic peaks. | 10 |
|  | DDA0017 | Relative standard deviations (%) to remove constant noisy peaks | 10 |
|  | DDA0018 | Pearson’s correlation coefficient threshold | 0.85 |
| **Creat unique spectra by measuring pairwise similarity across entire samples** | DDA0019 | Aggregation meta-variable (only when **DDA0005** is provided) | Name |
|  | DDA0020 | Export classified spectra plots for individual unique spectra | NO |
|  | DDA0021 | Mass window (Da) to match reference spreadsheet compounds and to detect similar DDA spectra | 0.01 |
|  | DDA0022 | Retention time window (min) to find DDA spectra variants or match reference spreadsheet compounds | 0.05 |
|  | DDA0023 | Minimum frequency (%) of detection for DDA spectra across all *.msp* files for untargeted DDA spectra | 10 |
|  | DDA0024 | Weighted spectra to measure entropy similarity score. Weighted functions are used to boost the intensity of low abundant peaks described by Li *et. al.,* **2021**, *Nature Methods* | TRUE |
|  | DDA0025 | Minimum spectral entropy similarity score to find similar DDA spectra | 0.9 |

**Table S.5.** Parameters for Spectra Library Search

| **Analysis step** | **Parameter ID** | **Parameter description** | **User provided input** |
| --- | --- | --- | --- |
| **MS/MS library annotation criteria on individual *.msp* files** | SPEC0001 | Number of parallel processing threads | 1 |
|  | SPEC0002 | Parallelization mode | Sample Mode |
|  | SPEC0004 | Precursor adduct types | NA |
|  | SPEC0005 | Mass accuracy to match precursor m/z (Da) | NA |
|  | SPEC0006 | Maximum retention time tolerance (min) | NA |
|  | SPEC0007 | Absolute spectral entropy difference (∆S) for library pre-filtering | 2 |
|  | SPEC0008 | Level of noise removal relative to the basepeak (%) similar to a method described by Li et. al., **2021**, *Nature Methods* | 1 |
|  | SPEC0009 | Minimum cutoff (%) for relative intensity ( = peak intensity/basepeak intensity) to select spectra marker peaks above this cutoff | 20 |
|  | SPEC0010 | Minimum percentage of the matched library and sample spectra markers (%) specified by **SPEC0009** | 50 |
|  | SPEC0011 | Level of pre-filtering which is a rounding digit to match rounded spectra markers for a quick pre-filter run. | 1 |
|  | SPEC0012 | Mass accuracy (Da) to integrate adjacent peaks and match fragments | 0.01 |
|  | SPEC0013 | Weighted spectra to measure entropy similarity score. Weighted functions are used to boost the intensity of low abundant peaks described by Li et. al., **2021**, *Nature Methods* | TRUE |
|  | SPEC0014 | Minimum distance (Da) between lowest and highest matched m/z to prevent matching only isotopic envelopes | 0 |
|  | SPEC0015 | Minimum number of matched peaks | 2 |
|  | SPEC0016 | Minimum spectral entropy similarity score | 0.75 |
|  | SPEC0017 | Minimum cosine similarity score | 0 |
|  | SPEC0018 | Maximum Normalized Euclidean Mass Error (NEME (mDa)) | 5 |
|  | SPEC0019 | Maximum number of top hits spectra figures for the plotting module | 0 |
|  | SPEC0020 | Spectra figure format | PNG |
